# Narrative ‘twist’ shifts within-individual neural representations of dissociable story features

**DOI:** 10.1101/2025.01.13.632631

**Authors:** Clara Sava-Segal, Clare Grall, Emily S. Finn

## Abstract

Given the same input, understanding of that input can differ depending on context. How does the brain represent the latent mental frameworks that support different interpretations of the same sensory information? In this study, participants listened to the same auditory narrative twice; the narrative had a plot twist in the middle that dramatically shifted interpretations of the story. Using a within-subject design that held both the stimulus and the individual constant, we leveraged reinterpretation-driven shifts in neural activity between the two listens to identify where representations of different narrative elements are updated under a new interpretative framework. We separated the narrative into three interrelated levels—the overall narrative model, episodes, and characters—to determine where reinterpretation-driven updates to each element were reflected in brain activity. Neural activity patterns associated with interpretations and reinterpretations of these three elements were observed in overlapping but partially distinct sets of temporal, parietal, and prefrontal regions. Results suggest that heteromodal cortex represents specific narrative elements according not to their surface features, but to latent conceptual frameworks for understanding those elements, which can shift under a new narrative interpretation.

**Significance statement:** Information is never experienced the same way twice. Here, we investigated where and how interpretations are updated in the brain when sensory input remains identical but contextual knowledge changes how we interpret that input. We used a unique narrative stimulus with a mid-story plot twist, which participants listened to twice. We found that sets of relatively dissociable regions across the brain represent three distinct narrative elements—narrative models, episodes, and characters. These findings reveal the neural basis for how identical information is processed differently through the mental frameworks we bring to understanding it.

## Introduction

Identical sensory inputs can evoke different interpretations. Rather than being fully predictable from properties of the information itself, our interpretations—how we understand or make meaning of external information—are flexibly shaped by how that information interacts with our internal expectations, prior knowledge, and mental state. Such flexibility is especially apparent in complex narratives, where construals of the situation, episodes, and characters can differ depending on one’s current interpretive framework.

Past neuroimaging studies have shown that varying interpretations of an identical narrative either by experimentally inducing opposing belief frameworks^1–3^ or by allowing participants to form their own endogenously generated interpretation^4–6^ evokes correspondingly distinct patterns of brain activity. Effects are frequently reported in regions of the default mode network (DMN), including the medial prefrontal cortex, posterior cingulate/precuneus, and angular gyrus. This is consistent with accounts that the DMN supports the construction and maintenance of high-level situation models, mentalizing, and schema-based predictions.

But what happens when we experience the *same* narrative under a *different* interpretive framework? While some prior work suggests that neural representations change as interpretations of the stimulus change^2,7^, many past studies have tracked narrative interpretations either by characterizing how neural representations evolve from the beginning to end of the narrative^8–10^ or by comparing representations between encoding and recall^2,11,12^. In both cases, differences in interpretations are confounded with changes in external sensory input, task demands, or mnemonic state, complicating efforts to pin differences in neural activity to differences in interpretive framework *per se*. Other studies have relied on explicit experimental manipulations to induce different interpretations of a given stimulus across subjects^1,3,13,14^. While informative, these studies cannot answer the question of if and how neural representations of the same information are updated under a *new* interpretive framework; furthermore, they somewhat compromise ecological validity since in the real world, we are rarely handed an explicit belief framework ahead of time. Here, we use a within-subject design in which interpretation and reinterpretation emerge spontaneously from the stimulus itself, allowing us to compare the same participant to themselves before and after a shift in interpretation. We thereby inherently control for any unwanted individual sources of variability (e.g., differences in prior experience, expectations, or brain functional anatomy) to examine not only where initial interpretations are instantiated, but also how a new interpretive framework reorganizes neural representations under largely naturalistic conditions.

A second open question is, as we form and change our understanding of a narrative, what exactly are we representing? Narratives are not monoliths, but are rather composed of separable elements that operate on distinct timescales. These elements include characters^10,12,15^ and episodes^2,8,10–12,16^ that dynamically interact with the broader narrative model. This narrative model (sometimes called a mental model or situation model^17,18^), which operates on a relatively long timescale, tracks the ‘gist’ of a story and provides the scaffold for interpreting these shorter-timescale elements, while also being reshaped by them. For example, when a character’s motivations are revealed or an unexpected event occurs, the overarching model is updated, which in turn alters how subsequent characters and episodes are understood. Behavioral evidence from prior work^19^ suggests that different narrative elements can be updated independently, raising the possibility that distinct neural systems may underlie the representation of specific narrative elements^12,20,21^. Prior work has mapped neural representations of narrative elements largely according to their surface features (e.g., the presence or absence of a given character or location^12^), which are stable regardless of interpretive framework, but not necessarily participants’ underlying conceptual understanding of those elements (e.g., a character’s attributes and motivations), which can shift under different interpretive frameworks. Other work has mapped where and how neural representations reflect deeper conceptual understanding of narrative elements, but has tended to limit focus to one element at a time (e.g., episodes as in^7,22^). Here, we broaden this scope to investigate where the brain tracks latent conceptual understanding of multiple narrative features simultaneously, focusing on three interrelated yet dissociable elements: narrative models, episodes, and characters. Moreover, rather than restricting analyses to a subset of regions chosen *a priori*, such as the default mode network (DMN) and related regions^2,7,10,12,15,23^, we take a whole-brain approach to understanding where and how interpretations of these elements are represented. The distinct timescales associated with each element predict that they may be represented across a hierarchy of brain regions with different temporal receptive windows for integrating information^8,24–26^.

In the current work, we provide a strong test of the idea that brain activity evoked by a given stimulus depends heavily on the interpretive lens through which it is experienced. Using repeated exposure to a unique narrative that contained a major twist halfway through, we hold both the participant and the stimulus constant, enabling us to leverage reinterpretation-driven changes in neural activity between the first and second listen to understand how and where representations of different narrative elements are updated. By modeling nested features of the narrative, we find that narrative models, episodes, and characters are represented in overlapping but dissociable sets of brain regions. Overall, using reinterpretation as a probe, we show that much of heteromodal cortex, including but not limited to regions of the default mode network, represents information not according to its surface sensory features, but according to latent mental frameworks for understanding that information.

## Results

Thirty-six healthy adults listened to an auditory narrative twice in a row during functional magnetic resonance imaging (fMRI) scanning. The format of the story is almost entirely a dialogue between a male and a female character, with some sparse additional effects (e.g., occasional background music) but no nondiegetic sound. The narrative features a twist in the middle that recontextualizes the earlier segments of the story: initially perceived as a straightforward interaction between a curmudgeonly dress-shopper (Steve) and a friendly, if pushy, shopkeeper (Lucy), the twist reveals a radically different reality: Steve is struggling to survive an apocalypse, and Lucy is a robot undermining his survival (See Supporting Info (SI), Methods section “*Stimulus description”* for further detail).

While repeated exposure affects how people process narratives^27,28^, unlike past designs that introduce the interpretation manipulation up front (*before* encoding; e.g.,^1–3,9^) or at the very end (*after* encoding; e.g.,^2^), here we introduce the manipulation *in the middle* of the first encoding experience, then have participants listen to the entire narrative a second time. This stimulus structure, whereby the ‘twist’ comes halfway through, triggers a relatively abrupt shift from one interpretation to another that lends itself to a straightforward hypothesis: interpretations, and therefore neural representations, of *pre-twist* events will change more than those of *post-twist* events. This allows us to control not only for sensory input, which is the same in both listens, but also for exposure: the entire narrative is experienced twice, but the most dramatic shifts should occur in the pre-twist period.

The shift induced by the twist required listeners to update their understanding of the broader narrative model, reevaluate specific episodes, and reassess the characters in light of the new context. We captured within-subject “*shifts*” – defined as between-listen changes in neural representations – to examine how reinterpretation-driven updates to neural activity patterns reveal distributed representations of each narrative element.

### Narrative elements are relatively dissociable

To examine how listeners represented different components of the story, we segmented the stimulus into three elements: *narrative model, episodes,* and *characters*, which operate at different timescales and capture interrelated yet distinct aspects of narrative information. First, the narrative model (longest timescale) captures the overarching situation, or ‘gist’, which gets reconfigured at the twist (i.e., shifting from “normal day at the mall” to “robot apocalypse”). Using *post hoc* participant reports (see SI Methods, “*Computing the ‘twist’”*), we divided the stimulus into three segments: pre-twist, twist, and post-twist (**Fig. 1, top row**). Second, we consider episodes (shortest timescale) as discrete plot-advancing events with a clear beginning and end. Participants indicated moments they reevaluated in light of the twist; we used these reports to identify five reevaluated episodes and matched nearby control episodes of identical duration (**Fig. 1, middle row**; see SI Methods, “*Computing shifts in the reevaluated episodes”*). Finally, we modeled characters (mid-level timescale) using the timestamps of speech turns taken by “Lucy” or “Steve” (**Fig. 1, bottom row**).

**Fig. 1.**
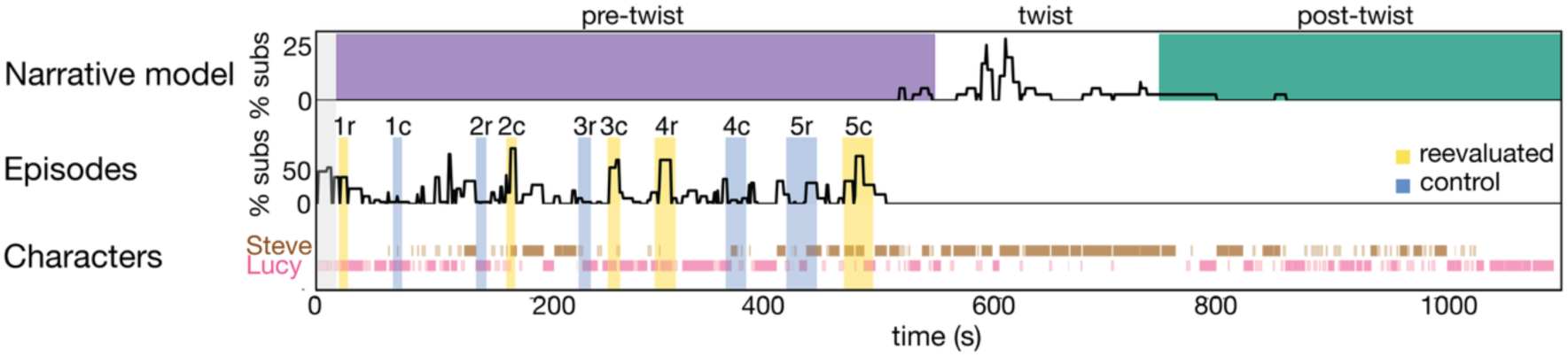
Timeline and dissociability of narrative elements. **Narrative model (top).** Black line: Time series showing, at each speech turn, the percentage of participants who marked that turn as the moment they first began to perceive the twist in *post hoc* behavioral reports. The pre-twist and post-twist segments as defined at the group level are indicated in purple and green. **Episodes (middle).** Black line: Time series of the percentage of participants who reported reevaluating each moment in *post hoc* behavioral reports (see Supporting Information (SI), Methods section, “*Computing shifts in the reevaluated episodes*”). Highlighted spans indicate the five reevaluated episodes (yellow) and their matched control episodes of similar duration and nearby timing (light blue; see SI, Methods), defined at the group level using these behavioral reports. Subscripts: *r,* reevaluated; *c*, control. **Characters (bottom).** Timestamps of speech turns for Steve (brown) and Lucy (pink). Low-opacity ticks denote very short turns (< 5 s) that were excluded from character-level analyses (Fig. 4). Leftmost gray vertical box corresponds to the first 18 seconds which were removed to avoid initial transient effects.

Although interdependent—episodes unfold through characters, and both contribute to the narrative model—according to our hypothesized framework, these elements capture distinct aspects of the story that might then be represented in distinct brain regions. Before modeling each element using the fMRI data, we therefore verified that they can be dissociated in terms of their structure and distribution.

Episodes typically spanned multiple character speech turns. We first verified that distributions of speaking time for both characters did not differ between the reevaluated and control episodes. Neither Lucy (Reevaluated:Control, M=62.79%±25.80%; M=65.78%±28.09%; t=-0.176, *p*=0.87) nor Steve (Reevaluated:Control, M=31.01%±29.57%; M=25.70%±18.36%; t=0.341, *p*=0.74) showed significant differences in their percent speaking time between reevaluated and control episodes. There were no differences in their proportion relative to one another (t=-0.113, *p*=0.913); nor in the variability of this balance across conditions (Levene’s test: *p*=0.51). This suggests that episodes and characters can be modeled independently.

For our character analyses, we verified that the two characters were generally balanced in their representation relative to one another and to the different segments of interest. (For these analyses, we limited models to speech turns that were at least 5 seconds in duration.) While Lucy spoke for slightly more of the narrative (40 speech turns versus 33 for Steve), the ratio of Lucy:Steve turns was consistent across segments of interest (pre-twist: 22:14 (∼1.6); post-twist: 16:9 (∼1.8), chi-square X^2^ = 0.052, *p* = 0.82). Turn durations were also comparable across segments: Lucy pre-twist mean = 10.32s (n=22), post-twist mean = 8.88s (n=16); Steve pre-twist mean = 8.93s (n=14), post-twist mean = 8.44s (n=9). Neither speaker showed significant differences in turn duration across segments (Lucy: t = 0.89, *p* = 0.37; Steve: t = 0.35, *p* = 0.73) nor were there differences in turn duration between speakers within segments (pre-twist: t = 0.85, *p* = 0.40; post-twist: t = 0.27, *p* = 0.79). Together, this suggests that shifts in the narrative model (operationalized as pre- versus post-twist segments) and shifts in understanding of characters (operationalized as speech turns) can be modeled independently.

The relationship between the narrative model and specific episodes is somewhat hierarchical. While all selected episodes (reevaluated and control) occurred in the pre-twist segment and thus underwent some amount of reinterpretation between listens, reevaluated episodes were moments when participants reported an especially strong shift in understanding given the information introduced by the twist. Comparing these reevaluated episodes to control episodes (**Fig. 3**) isolates moments where the updated narrative model should have its most dramatic effects on moment-to-moment understanding, providing a more sensitive test of interpretation updating.

The following results were robust to analytic choices, including adjusting the minimum duration of character turns or shifting boundaries (twist boundaries ±5–10 s, episodes ±3 s) to test reasonable alternative definitions based on the annotations.

### Representations of the narrative model

We first investigated where the overall narrative model—i.e., the broader gist of the narrative (in this case, “normal day at the mall” or “robot apocalypse”)—is represented and subsequently updated. To this end, we compared within-subject neural and behavioral responses between the two listens. Using participants’ *post hoc* behavioral reports of the moment they first began to perceive the twist, we split the narrative into three segments defined at the group level: pre-twist, twist, and post-twist. The twist changes the interpretation of everything that came before it, prompting participants to shift to a new narrative model that persists for the remainder of the first listen and throughout the second listen. As a result, the pre-twist segment, which is interpreted under a different model in the first (L1) versus second listen (L2), should be processed most differently between listens. In turn, the post-twist segment, which is processed with the same model across both listens, should exhibit more consistent neural and behavioral patterns (**Fig. 2A**). We therefore compared within-subject neural and behavioral shifts in each segment between listens (pre-twist_L1-L2_ to post-twist_L1-L2_), expecting greater shifts in the pre-twist segment than in the post-twist segment in both patterns of neural activity and behavioral ratings.

**Figure 2.**
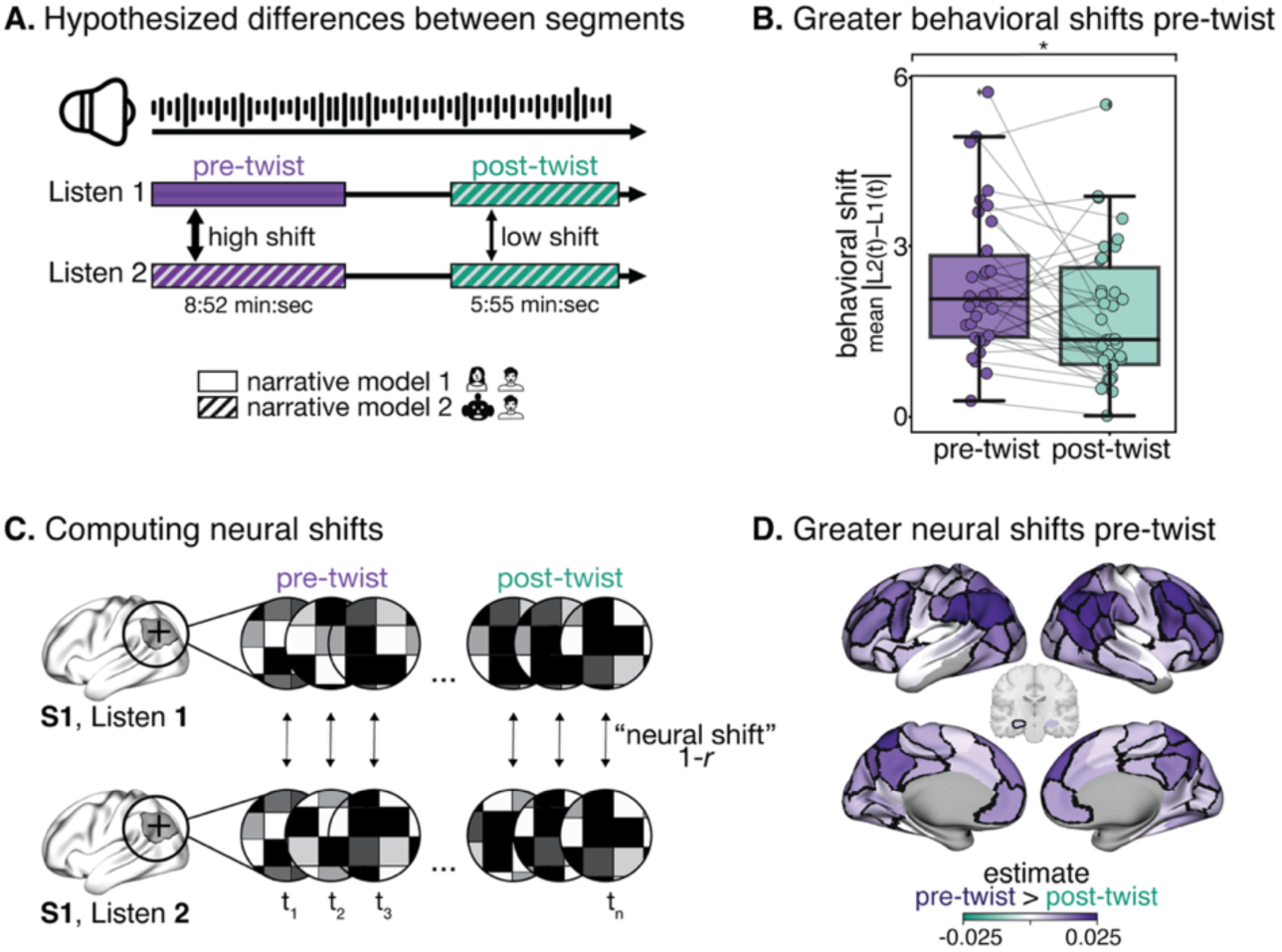
Neural and behavioral shifts reflect narrative model updating. **A. Hypothesized differences between segments.** The same individuals listened to an auditory narrative two times. The narrative was divided into three segments: 1. pre-twist, 2. twist and 3. post-twist. Novel, recontextualizing information is learned during the ‘twist’ segment, inducing a shift in interpretation. This new interpretation (narrative model 2) is carried into the post-twist segment on the first listen and into the entirety of the second listen. Thus, greater neural shifts are expected in the pre-twist as compared to the post-twist segment. **B. Greater behavioral shifts in the pre-twist segment.** Within-subject shifts between listens in behavioral ratings (continuous real-time reports of character impression) were greater in the pre-twist compared to the post-twist segment (paired t-test, * indicating *p* < 0.05). **C. Computing neural shifts.** For each participant, neural shifts between listens were computed per region per timepoint as one minus the correlation between the multivoxel spatial patterns of activity in Listen 1 and Listen 2 (pattern intra-SC). **D. Greater neural shifts in the pre-twist segment.** The median pre-twist and post-twist neural shift value for each participant was compared using a linear mixed effects model per region. Estimates plotted reflect the difference between the pre- and post-twist segments (set up as pre-twist > post-twist). Regions contoured in black show a significant effect at *q* < 0.05 (corrected for multiple comparisons using the false-discovery rate) for all matched-length sample comparisons between segments (see Supporting Information (SI), Methods).

#### Greater behavioral shifts in the pre-twist segment

During both listens, participants reported their real-time impression (negative to positive) of the shopkeeper (“Lucy”) in a continuous rating task. In the first listen, impressions of Lucy generally moved from positive to negative across the story, indicating a transition from viewing Lucy as a store clerk dealing with a difficult customer to an evil robot with Steve struggling to survive.

As hypothesized, within-participant ratings shifted more between listens in the pre-twist compared to the post-twist segment, indicating a greater change in how participants perceived the situation (“behavioral shift”; paired t-test, *t*(34)=2.36, *p* = 0.024, 95% CI [0.075, 1.005]; **Fig. 2B**). Behavioral shift was measured as the mean absolute difference between each participant’s Listen 1 and Listen 2 ratings within each segment.

Participants showed considerable variability in their compliance with this real-time behavioral task (see **Supplementary Fig. 1** for sample single-subject level behavioral reports). They consistently used a greater range of values, moved the slider more, and showed less predictable movement patterns in Listen 1 than in Listen 2. Further, this change in slider usage during Listen 2 manifested differently across participants, creating greater between-subject variability (including two participants that did not move the slider at all; see SI, Methods section – “*Variability in behavioral timeseries”*- for further detail). Therefore, while we initially had hypotheses linking shifts in behavioral ratings to shifts in neural activity on a finer-grained temporal scale, the combination of reduced task engagement and increased between-participant heterogeneity made it difficult to test these hypotheses reliably; we therefore did not use data from this task in any further analyses (aside from a control analysis described in the next section).

#### Greater neural shifts in the pre-twist segment

We operationalized neural representations as the multivoxel pattern of activity in each region at each timepoint. For each pair of matched timepoints in Listen 1 and Listen 2, we computed the within-subject correlation of these patterns (“pattern intra-subject correlation” [pattern intra-SC] ^29,30^) and calculated “neural shifts” as one minus this correlation (henceforth “intra-subject pattern distance”). As hypothesized (**Fig. 2A**), the intra-subject pattern distance was higher in the pre-twist segment than in the post-twist segment across the cortex, indicating greater neural shifts with reinterpretation (main effect of segment: estimate = 0.01, *p* < 0.001; whole-brain linear mixed effects model (LMEM) with region and participant as random effects). The regions that showed the strongest differences, suggesting a strong role in maintaining and updating the narrative model, included the left hippocampus, angular gyrus, temporo-parietal junction (TPJ), dorsomedial prefrontal cortex (PFC), and bilateral posterior medial cortex/precuneus (one LMEM per region with participant as a random effect; **Fig. 2D**). These regions have been implicated in across-subject studies on contextual modulation of representations of situation models and schemas ^12,16,31,32^ and in interpretational shifts during auditory narrative processing ^1,33,34^. They have also been characterized as integrating information over long temporal receptive windows ^24–26,34^. Together, this suggests that reinterpretation-driven updating of the narrative model preferentially engages cortical systems that accumulate and integrate information over extended spans of prior context.

Notably, we did not see significant neural shifts between listens in primary auditory cortex. This is an important negative control given that the low-level sensory properties of the stimulus are identical across listens. Some effects, albeit weaker than those in multimodal association regions, were also seen in early and middle visual regions (e.g., V1, MT). These effects are likely due at least in part to differences in how participants were looking at the screen to report or consider reporting Lucy impressions in the continuous rating task; in both listens, there were more slider movements (rating adjustments) in the pre-twist segment than in the post-twist segment (*t*(34) > 5.63, *p* < 0.001 for both listens).

We ran a series of control analyses to ensure the robustness of our findings. We first aimed to rule out the possibility that differences in brain activity across listens were driven by participants’ movements on the continuous rating task. To address this, we regressed the movement of the slider from each participant’s fMRI timeseries in each listen and repeated the analyses on the residuals of this regression; results were largely unchanged (**Supplementary Fig. 2A**).

Next, we sought to dissociate the effects of our stimulus’ specialized narrative model structure (i.e., the twist) from the effects of simply re-listening to the same information. First, one may expect that given that participants have already heard this narrative once, they may become less interested on the second listen. However, both our hypothesis and our observed results work against expected attention or “boredom” effects: if participants were simply mind-wandering more as time went on during the second listen, we would expect to see greater shifts post-twist compared to pre-twist due to decreased engagement and more off-task (as opposed to stimulus-driven) activity. A second possibility is that regardless of any narrative model updating, participants simply become more synchronized to themselves over time when relistening to the same narrative, which could also explain our pre- versus post-twist differences. We performed two analyses to help rule out this explanation. First, we turned to an independent dataset^35^ where the same participants listened to an auditory story (from *The Moth*) multiple times, of which we used the first two listens (**Supplementary Fig. 2B**). Critically, this story did *not* contain a twist or any other feature that would induce a drastic model update akin to our stimulus. Encouragingly, we found (at a liberal, uncorrected threshold of *p* < 0.05) that only three regions showed a linear effect of time on pattern intra-SC: dorsolateral PFC and the bilateral auditory cortex. The latter region (auditory cortex) actually showed a *decrease* over time, potentially indicating reduced attention. Second, in our dataset, we tested for any linear effects of time *within* segments (pre/post-twist) that could inflate our findings. We split the pre- and post-twist segments into an early and late period and identified where, regardless of segment (pre- versus post-twist), there were greater pattern intra-SC values in the late as compared to the early periods (**Supplementary Fig. 2C**).

Early versus late effects emerged in a small number of isolated regions (n=12), including the bilateral ventromedial PFC, the cingulate cortex, left superior STS (uncorrected *p* < 0.05) and the left intraparietal sulcus (*q* < 0.05). Only four of these regions (33%) showed effects in our primary analysis (see **Fig. 2D**), and effects in this control analysis were considerably weaker than those observed in the main contrasts. Results from this combination of analyses further strengthens the likelihood that our observed pre- versus post-twist differences were, in fact, driven largely by the narrative model updates induced by the twist in our stimulus, rather than simpler phenomena inherent to listening to the same stimulus a second time more generally.

### Representations of episodes

Having detected reinterpretation-driven updates associated with the overall narrative model in many regions of association cortex, we next investigated if and where the brain represents interpretations of more concrete units of a narrative, namely specific episodes ^36^. Here, we define episodes as punctate events with a clear beginning and end that advanced the plot (e.g., Steve trying on a wedding dress and Lucy remarking that he is skinny). Episodes are distinct from the larger narrative model not only in their timescale, but also in their level of concreteness: while they are more abstract than the text itself, they are still concrete in the sense that they are directly asserted by the narrative, unlike the global narrative model, which must be inferred by the reader and is therefore the most abstract^37,38^. Interpretations of particular episodes are revised under the new narrative model (e.g., recognizing that he is skinny because he is surviving an apocalypse). We hypothesized that, over and above the generally greater neural shifts in the pre-twist relative to post-twist segment, neural shifts in regions representing episode-level information should be even more exaggerated during the episodes whose interpretation shifted most dramatically under the new narrative model.

In a *post hoc* behavioral task, participants reported the episodes they reevaluated in light of the twist (see **Fig. 3A**; Supporting Information (SI) Methods section “*Computing shifts in the reevaluated episodes”*). We selected five episodes in the pre-twist segment that were reevaluated by the majority of participants along with five control episodes of matched length that were also in the pre-twist segment that most participants did not report reevaluating (shown in **Fig. 1**, middle row). Then, for each participant, we modeled patterns of brain activity during each individual episode in each listen using an event-related general linear model (GLM) and used the extracted episode-wise betas to compute a neural shift (intra-subject pattern distance between listens, **Fig. 3B**; see SI, Methods section “*Computing shifts in the reevaluated episodes”*). We further verified that these shifts were specific to the reevaluated episodes by comparing observed estimates to a null distribution of pseudo-episode pairs created by circle shifting (n=100) onset times within the pre-twist segment.

**Figure 3.**
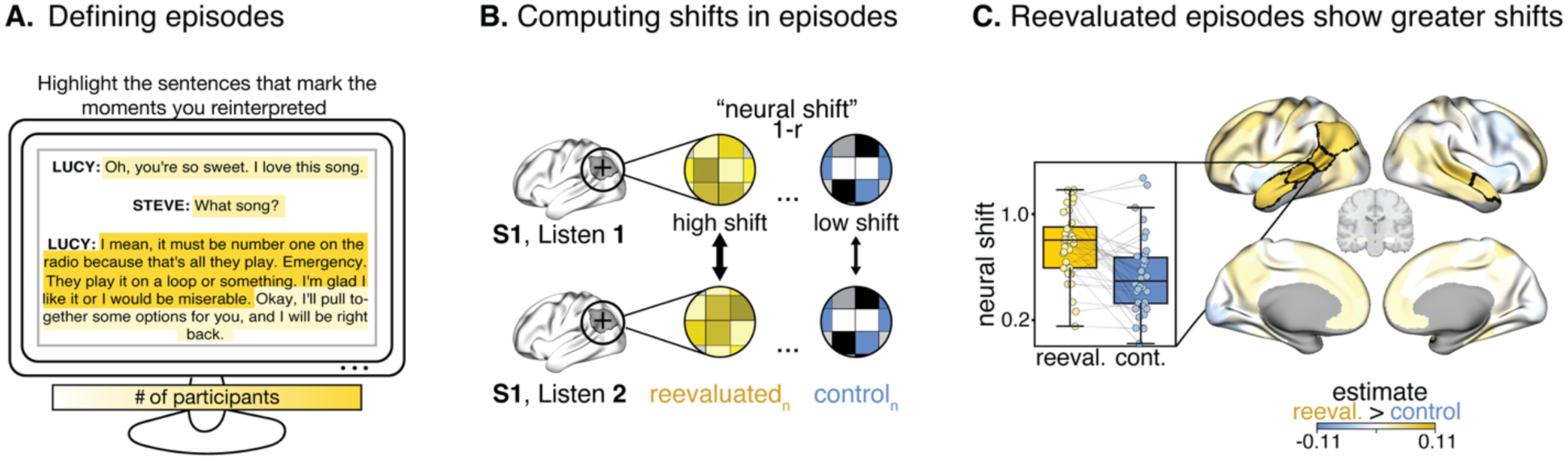
Episodes that are explicitly reevaluated show greater neural shifts between listens. **A. Identifying reevaluated episodes and matched controls.** After the scan, participants were given a transcript of the story and asked to highlight the moments where they experienced an especially strong shift in understanding in light of the twist; we used these highlighted transcripts as our primary source and cross-checked against spoken reports collected after each listen while participants were still in the scanner. We selected the top five episodes that participants most commonly reported reevaluating (**Fig. 1**) and paired each one with a matched control episode that was nearby in the narrative and the same length, but not reported as reevaluated by most participants. All reevaluated and control episodes were within the pre-twist segment. **B. Computing neural shifts between the reevaluated and control episodes.** For each participant, we used an event-related general linear model (GLM) to model each individual episode in each listen, then computed neural shifts as one minus the correlation between the spatial pattern of beta values in Listen 1 and Listen 2 (one value per region per episode). We hypothesized that neural shifts would be greater for the reevaluated compared to the control episodes. **C. Reevaluated episodes show greater neural shifts.** Plotted estimates show the strength of the difference between reevaluated and null episodes within participants. (Estimates reflect output from a linear mixed effects model in which within-subject neural shifts were predicted by episode type (set up as reevaluated > control), using participant and episode pair as random effects.) Regions contoured in black show a significant effect at *q* < 0.05 (corrected for multiple comparisons using the false-discovery rate). The distributions of neural shifts within the superior temporal sulcus are plotted in the inset. Dots represent participants’ median neural shifts (1 – Fisher z-transformed r-values) across episodes of each type (reevaluated and control).

By comparing neural shifts between reevaluated and control episodes, we found evidence that interpretations of episodes are represented along the bilateral superior temporal lobes and in the left temporo-parietal junction (TPJ; **Fig. 3C**; LMEM per region with both participant and episode pair [reevaluated, matched control] as random effects). These findings align with related work identifying the left anterior middle temporal gyrus and the TPJ (among other default-mode regions) as supporting ‘aha’ moments ^22,39^.

Compared to the broader narrative model, episode-level reinterpretation effects emerged in distinct, more left-lateralized temporal regions (compare **Fig. 2D** to **Fig. 3C**), which may be due to the left lateralization of language-mediated semantic representations ^40–43^. Notably, though, effects were strongest in the left superior temporal sulcus (STS) compared to neighboring regions more commonly associated with lower-level speech and language properties (i.e., auditory cortex and superior temporal gyrus; see SI, Methods section “*Comparing episode-level representations in the STS to neighboring regions”* and **Supplementary Fig. 3**). The one region that showed strong effects in both episode and narrative model representations was the left TPJ, a region suggested to be involved in binding of external information and managing competing beliefs ^44,45^. These distinct spatial patterns may also reflect differences in temporal receptive windows, as these sets of brain regions differ in their preferred information processing timescales. Specifically, episode-selective regions (superior temporal sulci/gyri) process shorter timescales while narrative model-selective regions (precuneus, mPFC) process longer timescales ^24–26^ with the TPJ as an intermediate.

### Representations of characters

Building on the reinterpretation-driven effects observed for narrative models and specific episodes, we next examined how representations of characters are constructed and updated. We investigated how representations of Lucy (from shopkeeper to robot) and Steve (from annoying shopper to persistent survivor) are constructed and updated across the two listens (see SI, Methods section “*Stimulus description”* for further detail on the characters).

We expected that by the end of Listen 1, participants would have converged on a final interpretation of the two characters and that they would then “reload” this interpretation into memory at the start of Listen 2. We operationalized these assumptions into predictions about what neural activity patterns should look like in regions tracking latent representations of the characters.

To this end, we split the narrative into “Lucy” or “Steve” speech turns based on speaking onset and offset times and modeled each turn in each listen using an event-related GLM (restricting the model to turns that were at least 5s long). We then designated a per-participant, per-region “template” neural representation of each character in Listen 1, when participants had all the information necessary to fully interpret (represent) their identity under the updated narrative model. Specifically, template representations were defined by taking the mean activity in each voxel across the final two speech turns of Listen 1 for a given character.

We then correlated representations of each character at each turn in Listen 1 and Listen 2 with their corresponding template, yielding a series of correlations per region showing how character representations evolve toward the template in each listen (**Fig. 4A**). Specifically, we considered a region as exhibiting reinterpretation-driven character-level representations if it met the following criteria: (1) a steady increase over time in similarity between the character’s speech turns and the template during Listen 1, (2) a reloaded template-like representation at the start of Listen 2, (3) a stabilization in representation (i.e., flatter slope toward the template) over the course of Listen 2, and (4) a dissociable representation of the other character (i.e., lower or negative correlations with the template that stay flat or decrease over time) in both listens (see **Fig. 4B** for a schematic of these criteria and **Supplementary Fig. 4A** for observed data from the left TPJ for both characters that show the hypothesized patterns). For more information on these criteria and how they were tested, see SI Methods section *“Computing updates in the representations of characters”*.

**Figure 4.**
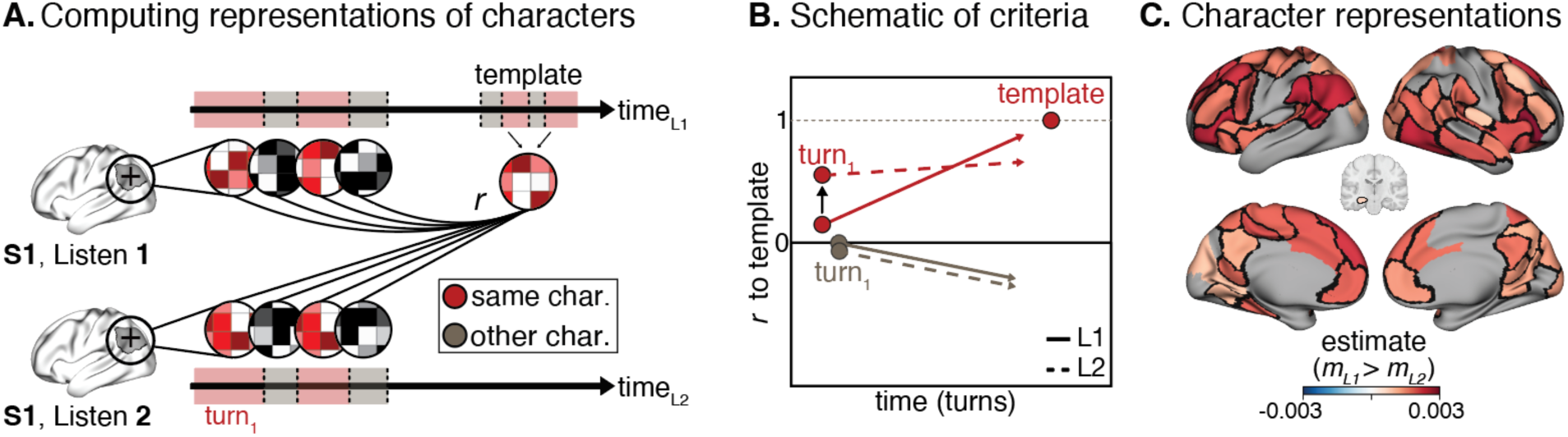
Character representations are updated in light of the twist. **A. Computing representations of characters.** The dialogue was split into ‘Lucy’ and ‘Steve’ speech turns based on speaking onset and offset times. We designated a per-participant, per-region ‘template’ representation of each character based on the multivoxel activity pattern during their last two speech turns in Listen 1 (L1). Multivoxel patterns during each turn in each listen were then correlated with the template to test for a series of criteria (see panel B). **B. Schematic of criteria.** Red lines: Pattern similarity (correlations) between same-character turns and the corresponding template (i.e., Lucy turns to Lucy template; Steve turns to Steve template) were hypothesized to be positive and to increase progressively over the course of the story (positive slope) in Listen 1. In Listen 2, they were expected to start higher and exhibit a weaker slope compared to Listen 1, reflecting the “loading” of the character’s representation from the end of Listen 1. Gray lines: Pattern similarity (correlations) between opposite-character turns and each template (i.e., Steve turns to Lucy template and vice versa) were hypothesized to be non-existent or, if anything, to show a negative slope over time (as representations of the characters diverged). Sample regions that show this hypothesized relationship are shown in **Supplementary Fig. 4A**. **C. Regions tracking character representations.** Estimates reflect the magnitude of the effect for our main criterion, which was that the similarity between a character’s turns and their template should show a steeper slope over time in Listen 1 than Listen 2 (computed with a linear mixed effects model predicting turn-template correlations from an interaction between listen and turn number with a random effect of participant). Regions plotted meet the thresholds for all of our criteria (see B; see Supporting Information, Methods). Black contours include regions that show a significant effect at *q* < 0.05 (following correction for multiple comparisons using the false-discovery rate) for the main criterion.

We focused on regions that showed effects in the expected direction for all criteria at an uncorrected threshold and that exhibited a significantly steeper slope over time (i.e., a sharper evolution toward the template) in Listen 1 relative to Listen 2, which we considered the most important criterion for a region representing latent character interpretations, at a corrected threshold (*q_FDR_* < 0.05; **Fig. 4C**). Regions meeting both of these standards emerged bilaterally in the posterior superior temporal sulcus (STS), posterior medial cortex (PMC), the temporo-parietal junction (TPJ), angular gyrus and ventromedial PFC for both characters ^10,12,20,46,47^. Effects were robust to choice of exact window size for template definition from 2-4 speech turns (see **Supplementary Fig. 4B**).

Taken together with the previous section, these results show that episode and character representations rely on somewhat distinct brain regions. Unlike episode representations, which are relatively localized to the left superior temporal lobe and TPJ, characters are represented bilaterally in more distributed medial regions, including posterior visual regions ^10,12^, parts of PMC, and mPFC).

### Dissociable neural substrates for representing distinct elements of narrative interpretation

Results thus far suggest that reinterpretation-driven neural representations of different narrative elements involve partially overlapping yet distinct sets of brain regions. This can be appreciated visually by comparing the maps for narrative models, episodes, and characters (compare **Fig. 2D** to **3C** to **4C**). To quantify this dissociation between the three narrative elements, we first assessed the degree of overlap in brain maps by correlating region-wise effect estimates across each pair of analyses. Regions encoding episodes were largely distinct from those encoding characters (*r* = 0.12, *n.s*.) or the narrative model (*r* = −0.02, *n.s*.). In contrast, regions encoding characters showed a weak but significant correlation with those encoding the narrative model (*r* = 0.24, *p* < 0.05), driven primarily by similarities in the spatial distribution of effect magnitudes within the default mode network (DMN; **Fig. 5A**), particularly within Cluster 1 (**Fig. 5B**).

**Figure 5.**
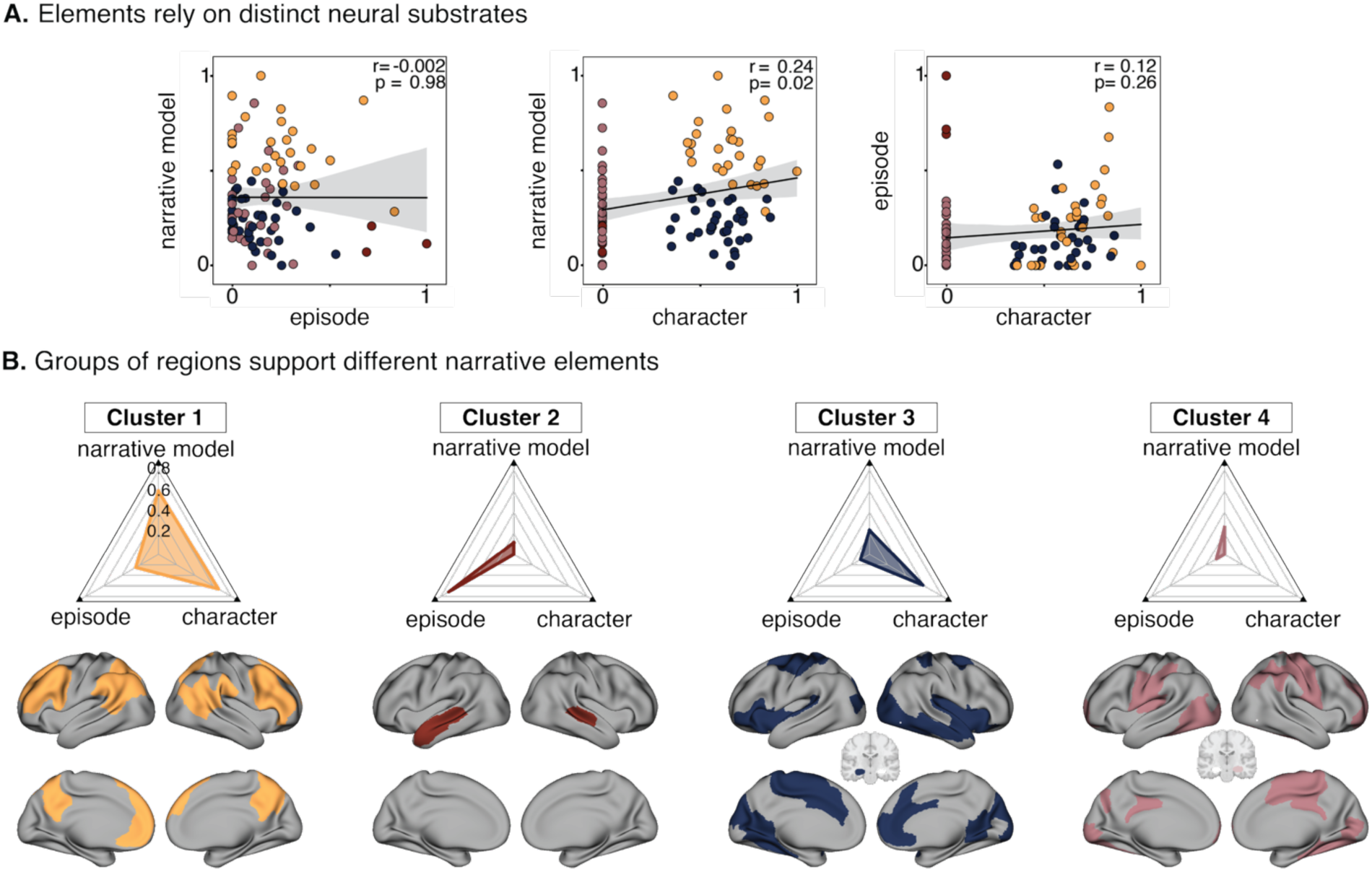
Representations of narrative elements are dissociable and linked to partially overlapping but distinct sets of brain regions. **A. Narrative elements are represented in distinct neural substrates.** Correlations between region-wise normalized effect estimates across each pair of analyses show small or nonexistent relationships, suggesting that representations of narrative models, episodes and characters rely on distinct neural substrates. Each dot indicates a region. Coloring of a dot (region) is based on the assigned cluster (see B). See SI Methods for further detail on the estimates that were set to 0. **B. Groups of regions support different narrative elements.** We clustered regions according to their relative involvement in representing the three narrative elements: narrative model, specific episodes, and characters. A solution of *k* = 4 clusters was found. Cluster 1 represented all three elements with a stronger weighting towards the narrative model and characters. Cluster 2 represented episodes and Cluster 3 represented characters most strongly while Cluster 4 showed weak involvement overall.

To further explore these distinctions, we applied KMeans clustering to group regions based on their distribution (pattern) of estimates from each analysis; this yielded a stable solution of four clusters (**Fig. 5B**). Results reinforce that the reinterpretation-driven effects linked to the three narrative elements are somewhat dissociable in where they are represented neurally.

To elaborate, the first three clusters had some involvement in representing a combination of the three narrative elements, but with varying relative weights. Cluster 1 represented all three elements, with a relative weighting toward narrative model and character representations. Notably, this cluster comprises the “core” regions of the default-mode network (DMN; bilateral angular gyrus, TPJ, precuneus/posterior medial cortex). These regions are thought to be involved in instantiating and updating broad contextual ‘gist’. Clusters 2 and 3 showed strong weighting towards episode and character representations, respectively, including the temporal regions often associated with the DMN. While Cluster 2 was restricted to the bilateral temporal sulcus and left temporal pole (regions thought to maintain representations of semantic information, identities, and mental simulations ^48,49^), Cluster 3 was more widely distributed including the right mPFC, right temporal pole and posterior medial visual regions. Cluster 4 (comprising some early visual and somatomotor regions) showed only weak involvement in narrative representations. Together, these analyses emphasize how reinterpretation drives updating in different sets of brain regions depending on which element is being updated.

## Discussion

In this study, we used reinterpretation as a probe to examine how latent representations of different narrative elements are updated in the brain. To do so, we deliberately selected an auditory narrative that featured a mid-story ‘twist’, or shift in the ground truth, that fundamentally altered participants’ understanding of earlier events. Participants listened to the stimulus twice, carrying forward the new narrative model induced by the twist into the second listen. This within-subject design enabled us to directly compare each participant to themselves as they updated their interpretations and understand how this interpretational shift altered the representation of the same sensory input. We decomposed the narrative into a general narrative model as well as two subcomponents, episodes and characters, and identified brain regions where multivariate activity patterns tracked not surface-level sensory features, but rather the conceptual understanding (interpretation) of these elements. We found that the broader narrative model exhibited the most widespread representations across the brain. In turn, episodes and characters engaged distinct sets of cortical regions, suggesting a degree of specialization in neural roles for representing and updating different narrative elements.

Much prior work has focused on the involvement of the default mode network (DMN) in narrative processing ^8,11,16,34,50^, namely showing that differences or changes in interpretations are reflected in differing activity in DMN regions ^2,4–7,10,26,51,52^. For instance, Zadbood *et al.,* (2022) used an effective across-subjects design and a movie with a plot twist at the end to demonstrate that representations in specific core subregions of the DMN (e.g., temporo-parietal junction [TPJ], medial PFC, temporal poles) varied based on participants’ prior knowledge of the twist during encoding *and* were updated with the new information gained via the twist during recall. Our findings align with and extend these prior across-subject studies. First, our use of a stimulus with a mid-story manipulation yielded extended periods under both a revised (pre-twist) and a stable interpretive framework (post-twist). This allowed for direct within-subject contrasts of encoding under different interpretive states. Second, we show that different regions and subnetworks of this larger network preferentially represent some narrative elements over others. While the core cortical DMN regions track the broader narrative model, lateral temporal regions such as the superior temporal sulcus (STS) and temporal pole appear to support more focal representations for episodes and characters. Taken together, our findings add to the longstanding evidence that the DMN comprises multiple, interacting subsystems with distinct functions ^53–55^.

These topographic distinctions may in part reflect known differences in how transmodal regions integrate information over time^9,16,26,56–58^. There is extensive evidence for a hierarchical processing architecture in the brain (see ^59^ for a review), in which early sensory regions track fast fluctuations in input, while higher-order transmodal regions integrate information over longer windows of prior stimulus context. In the present study, these distinctions emerged through reinterpretation-driven updating: regions with shorter receptive windows tracked more time-bound narrative elements such as episodes (e.g., left STS ^24,26,56^), whereas regions with longer intrinsic timescales showed stronger effects for broader narrative models (e.g., dorsolateral prefrontal cortex; see ^23,24,57^). Regions in lateral posterior parietal and temporal cortex, including and surrounding the TPJ, occupy an intermediate position along this hierarchy and were engaged in all narrative elements, consistent with a role in integrating and updating interpretations ^60^.

Importantly, the reinterpretation-driven approach here differs from the manipulations used in many previous studies of contextual modulation. Much prior work on contextual modulation has manipulated the *degree* of context for a given piece of information, varying the strength, amount, or immediacy of context while the underlying interpretive framework remains fixed (e.g., by presenting more or less preceding context for an incoming sentence ^9^ or by changing participants’ awareness of the causal relationship between events ^7,22^). In contrast, the present study manipulates the *nature* of the context: here, participants shift from interpreting events under one valid interpretive framework (i.e., Lucy-as-shopkeeper) to an equally valid, but different interpretive framework (i.e., Lucy-as-robot). Our findings therefore suggest that context-sensitive neural activity across different timescales tracks not only the presence versus absence of contextual support ^9,24,26,51^, but also nuanced differences in the implicit conceptual framework through which that information is interpreted.

There are several limitations to this work. First, our analyses rely on a single stimulus. Although this stimulus was carefully chosen for our study design, some observed effects may reflect idiosyncratic properties of this particular stimulus rather than more general features of narrative interpretation ^61^. This choice of stimulus led us to focus on two mid-level subcomponents of narratives—episodes and characters—that could be clearly defined in our narrative, though because the narrative was structured as a dialogue, the presence of a character was almost always tied to other narrative elements, making it more difficult to isolate characters in this particular stimulus than in prior work ^10,12,34^. Future research could extend to other narratives as well as broaden the scope to finer grained features, such as distinctions between main and secondary characters or hierarchical or nested event structures. Second, our mid-story twist and repeated-listening design enabled a strong within-subject approach, allowing us to compare matched segments that carried a new, shifted interpretation versus those that retained the original one. A complementary extension might include a control group that hears the twist before the first listen (but still experiences the narrative twice), providing a baseline in which the pre-twist segment is interpreted consistently under the same narrative model. Finally, the inclusion of an active task during and after narrative listening was a deliberate design choice, intended to capture real-time updating and to strike a balance between structured engagement and fully naturalistic listening. However, this task may have altered participants’ natural engagement with the narrative, potentially encouraging more deliberate or frequent updating of mental models than would occur during fully passive listening. In addition, variability in participants’ slider usage limited the use of this temporally precise behavioral measure of reinterpretation; future work could explore different approaches to stimulus annotation to help clarify the moment-to-moment relationship between conscious reinterpretation and shifts in neural representations.

In sum, we took a rigorous within-subject approach to capture how and where latent interpretive frameworks are instantiated and updated in the brain. By holding both the participant and the sensory input constant, we leveraged reinterpretation-driven shifts in patterns of neural activity to reveal how conceptual understanding of the same content at different levels—episodes, characters, and the overall narrative model—is represented differently under new context. Broadly, this work demonstrates how reinterpretation can serve as a tool for revealing the structure of conceptual representations underlying subjective experience.

## Methods Summary

Participants (n = 36) listened twice to an 18:25 episode of *The Truth* podcast (“Dark End of the Mall”), featuring a dialogue between Lucy and Steve, while undergoing functional MRI (fMRI; TR = 1 s; 3-mm isotropic voxels). The story includes a mid-narrative twist that prompts a global reinterpretation of earlier events. During scanning, participants continuously rated how much they liked Lucy (–3 to +3). After each listen (in the scanner), they verbally reported moments they reevaluated; after the second listen (outside the scanner), they highlighted transcript sentences marking the twist and the passages they reinterpreted (see Supporting Information (SI) Methods section “*Experimental procedures”*).

Following preprocessing (see SI Methods section “*fMRI data processing”*), we parcellated fMRI data into 97 regions (Schaefer 100 parcellation^62^ plus bilateral hippocampus^63^; five parcels excluded for missing data) for further analysis. We split the narrative into three distinct narrative segments (pre-twist, twist, and post-twist) and quantified within-subject “neural shifts” by correlating multivoxel patterns between listens, then modeling median shift per segment with linear mixed-effects models (LMEMs). See SI Methods sections “*Computing the ‘twist’” and “Computing neural shifts to assess narrative model representations”* for further details. We additionally ran a series of control analyses to ensure the robustness of our findings (see SI, Methods section “*Control analyses: ruling out possible confounding effects of time on within-subject similarity.”*)

To quantify shifts in the reevaluated episodes, we defined five reevaluated episodes and five control episodes that were matched in length and time (see SI Methods: “*Computing shifts in the reevaluated episodes.”*) Using a general linear model (GLM), for each participant, we modeled all episodes in each listen and then computed the correlation between the spatial pattern of the resulting voxelwise beta values in each region between Listen 1 and Listen 2. Lastly, per region, we fit a LMEM to test the hypothesis that neural shifts would be greater for the reevaluated compared to the control episodes.

To test where characters are represented, we segmented the stimulus into speaker-specific “speech turns”. For each participant and listen, we fit an event-related GLM with one regressor per turn to obtain voxelwise betas. For each character, a participant-specific template was defined as the median multivoxel beta pattern from the last two turns of Listen 1 (when the interpretation is finalized). For every non-template turn in both listens, we correlated its pattern with the corresponding template to test a series of four criteria that a region had to meet to be considered as representing character interpretations (described in detail in SI, Methods section “*Computing updates in the representations of characters.”* Additional checks were run to ensure the robustness of these findings, see SI.

To compare regional involvement across the three narrative elements, we (i) computed pairwise correlations of region-wise normalized effect estimates between analyses and (ii) grouped regions by their pattern of effect estimates using k-means clustering^64^. The number of clusters was chosen by the maximum silhouette score (k=4). See SI, Methods “*Clustering regions across analyses”* for further details on these procedures.

## Code availability

Data analysis, including links to code and other supporting materials, can be found at: https://github.com/thefinnlab/darkend_narrative_rep.

## Data availability

Data from this study, including raw MRI data, will be made available on OpenNeuro upon publication.

## Acknowledgements

The authors thank Josefa M. Equita for assistance with data collection and behavioral preprocessing; Eneko Uruñuela for implementing denoising with Tedana; Katherine Bartolino and Evan Bloch for their contribution to behavioral data cleaning and processing; Rekha Varrier for early discussions on study design and data collection; Thomas L. Botch for preprocessing the *Moth Stories* data and providing thoughtful comments on the manuscript.

