## Supplementary figures for "Narrative ‘twist’ shifts within-individual neural representations of dissociable story features"

### Methods

#### Stimulus description.

We used the “Dark End of the Mall” episode (18:25 min:sec) from the podcast *The Truth*, which consists almost entirely of a dialogue between two characters, Lucy and Steve [1]. There is no non-diegetic sound, and the non-speaking time is limited to moments of a dog barking and brief moments of a “song” playing in the background. We chose this stimulus due to the feasibility of working with only two characters and, more importantly, its unique narrative structure. Specifically, the narrative contained a twist in the middle that required participants to globally update their narrative model of events that preceded the twist, creating three distinct and meaningful narrative segments (pre-twist, twist, and post-twist) for within-subject comparison. We provide a brief synopsis below.

> *The story starts off with a phone conversation between Lucy, a sweet but vapid bridal shop employee, and presumably her boyfriend (whom listeners do not hear) which gets interrupted by Steve running into the shop. Listeners initially perceive Steve as a cranky dress shopper who is abrasive toward Lucy as he tries multiple attempts to convince her that she should give him some of the food hidden in the shop. Lucy gets frustrated with Steve and calls mall security and tries to kick him out of the shop. Eventually, Steve asks Lucy if he can tell her a story. It is revealed via Steve’s story that Lucy is, in fact, a robot programmed to work in a 1950s style bridal shop, that they are both living in an apocalypse in 2050, and that Steve is one of the last surviving humans and has figured out that bridal shops have hidden snacks that sustain his survival. He almost convinces Lucy to help him, but ultimately fails as she kicks him out of the shop where, presumably, he meets his death. Listeners last hear a distressed Steve confronting barking dogs and Lucy again on the phone with her boyfriend, but listeners now realize via the narrative that the dogs are likely zombies and the boyfriend is fictitious.*

**Participants.** All data was collected at the Dartmouth Brain Imaging Center. Participants (n=36; 11M; median age = 20, range = 18 to 33) were healthy individuals, with normal or corrected-to-normal vision and hearing and no recent psychiatric or neurological diagnoses or MRI contraindications. They were recruited from the local areas of New Hampshire and Vermont, including the Dartmouth College student body. The Committee for the Protection of Human Subjects of Dartmouth College approved the study, and all participants provided written consent.

**Experimental procedures.** All participants listened to the same auditory stimulus twice. At the beginning of the study, they were told that they *may* hear the auditory narrative a second time, but that they also may have the opportunity to hear a second story. No participant actually heard a different story. While in the scanner, we used Sensimetrics Model S14 insert earphones to present the sound, and participants were given a trackpad (Cedrus Lumina) to continuously indicate their impressions of Lucy from very negative to very positive throughout each listen. They were given minimal visual input: the screen displayed throughout both listens showed a static photograph of a bridal shop (to promote imagery and engagement with the story) and, underneath that image, the continuous scale used to rate Lucy impressions.

*Continuous Rating Task.*

Participants were tasked with rating the character of Lucy by answering the question: “*Overall, how much do you like Lucy?*” During the presentation of the stimulus, participants used the trackpad to update their rating along a scale from -3 (very negative) to 3 (very positive) while the stimulus played. We opted to do this task in real time in the scanner as opposed to in an independent dataset of non-fMRI participants because pilot participants showed considerable variability in their ratings. Furthermore, to emphasize the within-subject design of our study, we did not want to use other participants’ data as a proxy for fMRI participants’ ratings of the character.

Before the second listen, participants were instructed as follows: "*For your 2nd story, you have been assigned to listen to the same story again and complete the same prompt. For this 2nd listen of the same story, consider how your impression has changed. Because you have already listened to the story, we expect that your impressions of Lucy are different than your 1st listen. Given what you know about this story, what is your impression of Lucy now? Please use the slider to continuously rate your impression.*"

*Tasks after each listen.*
Each scanner run consisted of the entire narrative; after each listen, participants were asked to do a series of character rating questions and memory tests (maximum of 10 seconds each) and engage in a "reevaluation task". For the character rating questions, participants were asked to report "Overall, how much do you like [Lucy/Steve]" as independent questions. For memory questions, participants were asked "*1. What is Lucy?, 2. What does Lucy hear running throughout the story?, 3. What is the name of the shop where this story takes place?"* after Listen 1 and "*1. What caused the destruction of humankind?, 2. What dish does Lucy recommend Steve buy at the food court?, 3. Who does Lucy think she is talking to at the beginning of the story?"* after Listen 2. These questions were intended to be relatively challenging and therefore to serve as attention checks. Participants performed well on these questions (median score 100%; mean score = 93%). After completing these questions, participants then completed the reevaluation task after each listen. After Listen 1, they were instructed "*Using the microphone, please describe the moments at the beginning of the story that you reconsidered after hearing the end."* After Listen 2, they were instructed, "*please describe the moments of the story that changed for you after hearing the story once before.*" They had 60 seconds to answer using free speech. Sample transcripts from three subjects are included in Table 1 in the Supporting Information.

*Post-scan tasks.*
Outside the scanner, participants were presented with the transcript and asked to "*highlight the 1-3 sentences that mark the moment in the story when the twist occurred."* They were also given all of the sentences in the pre-twist segment and tasked to indicate the ones that they reevaluated. Instructions stated "*highlight the sentences that mark the moments in the story that you reinterpreted when listening to it a second time*."

**fMRI data processing.**
*MRI acquisition.* All data were collected at Dartmouth College in a 3.0 Tesla Siemens MAGNETOM Prisma whole-body MRI system (Siemens Medical Solutions, Erlangen, Germany) equipped with a 64-channel head coil.

*T1 image.*
For registration purposes, a high-resolution T1-weighted magnetization-prepared rapid acquisition gradient echo (MPRAGE) imaging sequence was acquired (TR = 2,300 ms, echo time (TE) = 2.32 ms, inversion time = 933 ms, flip angle = 8°, field of view = 256 × 256 mm, slices = 255, voxel size = 3 × 3 × 3 mm isotropic). T1 images were segmented, and surfaces were generated using FreeSurfer [2].

*fMRI acquisition.*
fMRI data were acquired using a multi-echo T2*-weighted sequence. The sequence parameters were: TR = 1,000 ms, TEs = [14.2, 34.84, 55.48], GRAPPA factor = 4, flip angle = 60°, matrix size = 90 × 72, slices = 52, multiband factor = 4, voxel size = 3mm isotropic. To account for field stabilization and hemodynamic delay, an additional two seconds were added to the front of the stimulus and 10 seconds were added to the end.

*Preprocessing.*
Multi-echo data preprocessing was implemented in AFNI [3] using afni_proc.py for alignment, transformation, and optimization steps. Each participant's data was processed to align the anatomical (T1) image and functional images, with motion correction based on the second echo and alignment parameters applied to all echoes. Functional data underwent despiking (3dDespike)

for outlier attenuation, followed by the concatenation and extraction of functional time series for each echo. The three echoes were then optimally combined and denoised using multi-echo ICA via *tedana* [4–6]. Signals were then normalized to percent signal change and spatially blurred (3dBlurInMask), with motion regressors applied to reduce artifacts in final volumes. Following preprocessing, to account for transitory changes at the start of the stimulus [7], we removed the first 18 seconds from the start of the stimulus for all of our subsequent analyses (grayed out in **Fig. 1**).

*Defining regions of interest.*
The Schaefer parcellation[8] was used to designate 100 cortical regions; five of these regions—around the ventral part of the brain—were removed because more than 50% of participants were missing more than 40% of the data in these regions. The Harvard-Oxford Atlas was used to identify the hippocampus in both the left and right hemispheres[9]. We were unable to include other subcortical regions, including the amygdala, due to data loss (almost 50% of participants (17/36) were missing signal in more than 40% of voxels). Parcel sizes ranged from 113 to 759 voxels. All results shown here were robust to parcellation granularity in that effects persisted when using a 400-region parcellation[8].

**Computing the 'twist'.**
We defined the "twist" in the story as moment(s) when participants transition from one interpretation/narrative model to another—specifically, from believing the setting is a bridal shop to realizing it is a post-apocalyptic world. To capture this shift, participants were asked to identify the twist in a post-scan survey (see *Experimental procedures* for more information on instructions). Participant responses varied considerably, with some selecting multiple points in the story. These responses are shown in **Fig. 1.** To address this variability, we adopted a conservative approach to identifying the twist, defining its start as the point before the earliest event chosen by the majority of participants, and its end as the point after the latest event chosen by the majority. This approach allowed us to split the stimulus into pre-twist (length = 532 s) and post-twist segments (length = 355 s), plus a segment in the middle corresponding to the twist itself (length = 200 s). Pre- and post-twist segments were matched for length when appropriate (see Section Methods: "*Computing neural shifts to assess narrative model representations"*).

**Computing behavioral shifts to assess narrative model representations.**
We compared "behavioral shifts" between the pre-twist and post-twist segments using the timeseries from each participant's continuous rating of Lucy acquired in both listens. See *Experimental procedures* for specific instructions on how this continuous rating task was conducted. To measure behavioral shift, we calculated the absolute difference between each participant's Listen 1 and Listen 2 continuous ratings at each timepoint, then averaged these differences within each segment. This yielded a single behavioral shift score per participant per segment, with higher values indicating greater change between listening sessions. We compared these scores between pre-twist and post-twist segments using paired t-tests. Two participants did not move the slider at all in the second listen, yielding n=34 for this analysis.

**Variability in behavioral timeseries.**
Within individual participants, slider usage varied significantly across listens. Participants made more slider adjustments in Listen 1 than Listen 2 (Md=20.5 vs 16.0; Wilcoxon signed-rank test, $p$ = 0.044), used a greater range of values (M=5.88±1.53 vs 3.91±1.98; paired t-test, $p$ < 0.001), and had lower movement entropy in Listen 2, indicating more predictable patterns (M=0.163±0.104 vs 0.142±0.135; Wilcoxon signed-rank test, $p$ = 0.031). Between-participant variability was higher in Listen 2 across slider metrics (range SD: 1.53 vs 1.98; entropy SD: 0.104 vs 0.135; Levene's test for equal variances approaching significance, p=0.053). This inconsistency reduced the sensitivity of the behavioral timeseries, limiting their utility for reliable neural–behavioral analyses. Therefore, we did not try to find relationships between behavioral shifts and the neural data, discussed below.

**Computing neural shifts to assess narrative model representations.**

Our first goal was to quantify changes in the within-subject representation of the narrative ("neural shifts") between listens and to compare the magnitude of these changes between the pre-twist and post-twist segments. For each participant and region, we correlated the multivoxel spatial pattern at each timepoint between listens, yielding a pattern intra-subject correlation (intraSC)[10,11] for each timepoint. We Fisher z-transformed these correlations and then converted them into a "neural shift" (i.e., distance) at each timepoint by subtracting the pattern intraSC from one.

To test for a difference between segments, we computed the median neural shift value within each segment for each participant and conducted a linear mixed effects model (LMEM; using lme4 in R; [12]) where median neural shift was predicted by segment (pre- or post-twist) with participant as a random effect. Note that taking the median neural shift from each segment, as opposed to using shifts from all timepoints, helps accounts for autocorrelation in the functional data which would otherwise artificially inflate degrees of freedom. One LMEM was run per region. Estimates from each of these LMEMs are plotted in **Fig. 2D**.

To ensure observed neural shifts were not driven by differences in length of the two segments (8:52 min:sec versus 5:55 min:sec, or 532 versus 355 TRs), we trimmed the pre-twist segment to match the length of the post-twist segment. Specifically, we generated all possible 355-TR subsets of the pre-twist segment by sequentially trimming the pre-twist data from the start, creating 178 distinct samples (532 - 355 + 1). For each sample, we ran an LMEM for each region to compare the pre-twist and post-twist segments. The p-values from these models were then corrected for multiple comparisons using false discovery rate (FDR) based on the number of regions in our analyses (97 total: 95 cortical and two hippocampi) using an alpha of 0.05. To be as conservative as possible, we only considered regions to be significant if they were $q_{FDR} < 0.05$ in all 178 matched-length samples (**Fig. 2D**).

**Control analyses: ruling out possible confounding effects of time on within-subject similarity.**
To further ensure that the observed results, which were consistent with our hypothesized directionality (higher similarity in the post-twist relative to pre-twist segment), were due primarily to shifts in interpretation rather than other explanations, we investigated the alternative hypothesis that individuals simply become more similar to themselves over time when processing the same long-timescale narrative. Critically, we tested this hypothesis both in an independent dataset as well as in our own dataset, as described below.

*Computing within-subject similarity over time in an independent dataset.*
We used fMRI data from an existing dataset [13] in which participants (n = 8) listened to the same *The Moth* story ("*Where There's Smoke*") multiple times. Importantly, this story lacked a twist or any other feature that might induce interpretational differences, making it suitable as a control. We used data from the first two times participants listened to the story (run 1 from session 2 and run 2 from session 3), performed functional alignment using hyperalignment [14] with a leave-one-session-out cross-validation procedure, and again parcellated the data using the Schaefer parcellation. Taking the same approach as in our main analyses, we then computed the pattern intra-SC at each timepoint. To assess whether within-subject similarity changed with time, for each region, we fit a linear model for each participant predicting pattern intra-SC as a function of timepoint (TR). We then evaluated statistical significance using a one-sample t-test (two-sided) for each region on the resulting beta values across participants. We applied a liberal, uncorrected threshold of $p < 0.05$ to see which regions, if any, showed increased or decreased similarity over time (**Supplementary Fig. 2B**).

*Comparing early versus late changes within segments of our narrative.*
To further investigate possible confounding effects of time (rather than interpretation) on the similarity of within-subject neural representations, in our dataset, we further divided our pre-twist and post-twist segments into early and late halves. This resulted in four distinct periods: pre-twist early, pre-twist late, post-twist early, and post-twist late. Within each of these four periods, we took an approach mirroring our main analysis: for each participant, we computed pattern intra-SC values

between listens at each timepoint within each region. We then tested the hypothesis that intra-SC (similarity) would be higher in the late compared to early periods of a segment within each segment type. To do so, we fit a linear mixed effects model per parcel predicting median intra-SC from segment type (pre-twist versus post-twist) and half (early versus late), with random effects of participant. Effects are plotted in **Supplementary Fig. 2C**.

**Computing shifts in the reevaluated episodes.**
As discussed in further detail in *Experimental procedures,* after each listen, participants reported episodes – distinct events with a clear beginning, middle, and end – that they reevaluated. Specific reevaluated episodes were identified for analysis using two complementary data sources. During an in-scanner task, participants verbally described moments that had changed in meaning after hearing the complete narrative. These responses were transcribed and matched to specific story segments. Participants were quite variable in the number of and detail associated with episodes they reported reevaluating; sample responses can be found in **Supplementary Table 1.** Subsequently, participants completed a post-scan computerized survey highlighting sentences in the story transcript that they had reevaluated ("*Post-scan tasks*"). The post-scan data served as the primary source because identical text enabled more direct cross-subject comparison than varied verbal descriptions. A distribution of reevaluation frequency was computed across each second of the story to identify episodes consistently reported by participants. These were then manually checked for convergence with the in-scanner verbal report data. The highlighting data is made available in the code repository.

This process led us to select five episodes consistently noted by the majority (at least 25/36; ~70%) of participants in both their in-scanner verbal reports and post-scanner written highlighting task. These episodes varied in duration (8, 9, 11, 18, and 27 seconds) and were manually checked by the experimenter to ensure that they included the entirety of an episode, i.e., if a participant chose only one of the two sentences that comprised an episode, we considered the entirety of the episode if the majority of other participants reported reevaluating all of it. Importantly, all identified episodes occurred within the pre-twist segment (**Fig. 1**).

We then selected a set of "control episodes" to serve as a comparison point for the reevaluated episodes. To this end, we identified episodes within the pre-twist segment that the majority of participants (no more than 11/36; less than 30%) did *not* report reevaluating. We intentionally chose these episodes to be matched in length and nearby in time to the reevaluated episodes to account for any signal drift and to ensure that both the "control" and "reevaluated" episodes were within the same pre-twist segment (which had more overall reinterpretation, see **Fig. 1**). A brief description of these episodes is provided below.

> *The reevaluated episodes include the following: 1r. a conversation that Lucy has with an imaginary boyfriend; 2r. when Steve calls her a robot (reinterpreted from 'corporate drone' to actual robot); 3r. when Lucy calls Steve skinny which participants begin to realize is because he has been in survival mode for years; 4r. the 'emergency song' which is not a 'hit song' of the summer, but rather an emergency signal in the apocalypse; 5r. when Lucy tells Steve that the reason she cannot give him food and water is policy (because she is programmed to prevent this). The corresponding 'control' moments are when 1c. Lucy welcomes Steve to the store; 2c. when Lucy is impressed with Steve's knowledge of the store's policies; 3c. when he asks if his trying on a dress is against their policy; 4c. when Lucy chastises Steve; 5c. and when he calls her kind.*

To compute the timings of each episode, we first used WhisperX [15] to force-align the stimulus transcript with the auditory narrative. This process yielded an onset and offset timing for each word in seconds. We defined each episode as lasting from the onset of the first word to the offset of the last word. It is important to note that the start of the first reevaluated episode was excluded because it overlapped with the portion of the stimulus excluded to account for transitory delays (i.e., therefore, we only included the remaining portion of the episode).

Next, to directly assess whether the reevaluated episodes showed greater processing differences between listens compared to control episodes, we applied a general linear model (GLM) analysis. This pattern-level approach to episode updating is similar to that taken in prior work [16,17]. Using a GLM, for each participant, we modeled all episodes (10 total; reevaluated and control) in each listen using individual regressors for each episode (implemented as an individual-modulated event-related analysis using AFNI's *3dDeconvolve* function). This allowed us to obtain voxel-wise beta values for each episode. We then computed the correlation between the spatial pattern of voxelwise beta values in each region between Listen 1 and Listen 2, Fisher z-transformed these correlations, and calculated 'neural shifts' for each episode as one minus this transformed coefficient (**Fig. 3B**). Lastly, per region, we fit a LMEM to test the hypothesis that neural shifts would be greater for the reevaluated compared to the control episodes. This was set up using a main effect of episode type (set as a contrast of reevaluated > control), using a random effect of participant and episode pair. Here, episode pair refers to each pair of reevaluated and control episodes that were close in time and matched in length. P-values from the models were corrected for multiple comparisons using FDR with an alpha of 0.05, based on the number of regions analyzed (97; **Fig. 3C**).

To further validate resulting effects, we performed a permutation test by circularly shifting the onset of the reevaluated and control episodes within the pre-twist segment by a random amount across 100 iterations. We then applied the same GLM and statistical modeling approach to each permutation, generating a null distribution of parameter estimates for each brain region. We then compared the observed estimates from regions showing significant effects in our main analysis ($q_{FDR} < 0.05$, **Fig. 3C)** against their respective null distributions. All regions that showed significant effects in the main analysis remained significant in the permutation test ($p < 0.05$).

**Comparing episode-level representations in the STS to neighboring regions.**
Some of the strongest episode-related effects were observed in the left superior temporal sulcus (STS). Because STS is adjacent to regions more strongly associated with auditory and linguistic processing (the superior temporal gyrus [STG] and auditory cortex), we tested whether these effects reflected narrative reinterpretation specifically rather than more general auditory/semantic processing. To do so, we compared neural shifts for reevaluated versus control episodes in the left STS with those in neighboring temporal lobe regions, including subdivisions of the superior temporal gyrus (the planum temporale and lateral STG) and the primary auditory cortex (the transverse temporal sulcus and Heschl's gyrus), using the anatomically defined Destrieux cortical parcellation, which explicitly distinguishes sulci from gyri[18]. We applied the same analytic approach as in the previous section to compare reevaluated and control episodes within each of the six regions. To assess between-region differences in the magnitude of neural shifts, we compared participant-level neural shift estimates across regions using paired t-tests. P-values were corrected for multiple comparisons using false discovery rate (FDR) correction across the six regions analyzed. A visualization of the outputs can be found in **Supplementary Fig. 3.**

**Computing updates in the representations of characters.**
In our final analysis, we aimed to track where and how characters are represented. To this end, we segmented the stimulus into blocks in which either Lucy or Steve was speaking (speech "turns"). We identified the onset and offset of each turn using the word-level alignment times provided by WhisperX [15] and manually verified. We excluded turns shorter than 5 seconds (such as Steve saying his name), resulting in 40 turns for Lucy and 33 turns for Steve (median turn length: 9 seconds; range 5-27 seconds).

For each participant, we ran a GLM for each listen with individual regressors for each speech turn (implemented as an individual-modulated event-related analysis using AFNI's *3dDeconvolve* function, similar to the episodes analysis described above). Then, within each region, we took the mean across voxel-wise beta values from the last two character-specific turns of Listen 1 to define a "template" representation for each character (i.e., last two Lucy and last two Steve turns). Specifically, in these turns, participants should have a finalized understanding of who each character is under their revised interpretation following the twist.

Thus, for each participant, we correlated the multivoxel patterns of beta values for each non-template turn in both Listen 1 and Listen 2 (either Lucy or Steve) to the participant's own character template (**Fig. 4A**) and Fisher z-transformed these correlations. Lastly, we leveraged these turn-template transformed pattern coefficient values to identify character representations (tested using the following criteria).

***Criteria for character representation updating.***
We expected that representations of each character to evolve for participants throughout Listen 1 (e.g. for Lucy, from shopkeeper to robot), and that the updated representation would be "loaded" back into memory at the start of Listen 2. We evaluated which brain regions exhibited this representational transition—and could therefore be considered to instantiate latent interpretations of characters—as defined by the following criteria. In all of the following tests, data from both characters (Steve and Lucy) were analyzed in the same LMEMs with random effects of participant and character, to account for between-participant and between-character variability.

*Criterion 1— Representations of the characters become more like the template throughout Listen 1.*
To test if representations of a character become more like their respective template, we fit a LMEM for each region, predicting the speech turn-template Fisher z–transformed correlation in Listen 1 from the turn number (with higher numbers corresponding to later turns) and treating participant and character as a random effect. We hypothesized a positive linear trend across character speech turns, with later turns showing stronger correlations with the template as representations converge toward the final template, reflecting participants' learning about the character across the first listen. For this criterion to be met, the statistic had to be positive and significant at an uncorrected threshold $p < 0.05$.

*Criterion 2—Representations of the characters during their first speech turn are 'updated' in Listen 2.*
To test if participants "load in" their updated representation of a character when starting Listen 2, we compared the correlation (Fisher z–transformed) to the respective template for the first character speech turn between the two listens. By comparing the same character speech turn (matched sensory input) across listens to the same template, we considered that stronger correlations with the template in Listen 2 reflected a shift toward a more updated representation of that character.

For each region, we fit two LMEMs that pooled data from both characters and included random effects for participant and character. To assess whether correlations between the first speech turn and the template increased from Listen 1 to Listen 2, we fit a model with Listen as a fixed effect with a contrast comparing them (Listen 2 > Listen 1). We additionally tested whether correlations in Listen 2 were reliably greater than 0 by fitting an intercept-only model to the Listen 2 data. For this criterion to be met, both statistics had to be positive (i.e., Listen 2 > Listen 1; Listen 2 > 0).

*Criterion 3—Representations of characters stabilize in Listen 2.*
To test if representations of either character "stabilize," we compared how speech turn-template correlations evolved over time between the two listens. We fit a LMEM for each region, predicting the turn-template z-transformed correlation based on an interaction of listen (Listen 1 or Listen 2) and character speech turn number, treating participant and the character as a random effect. As noted in Criterion 1, we hypothesized that there would be a positive linear fit of turn number—that is, later character speech turns would be more correlated to the respective template as the representation built up over the course of the narrative. Here, additionally, we tested that the positive slope would be steeper in Listen 1 relative to Listen 2, given that in Listen 2 the character representation requires less updating and starts out closer to the template because it has been "preloaded" into memory. For this criterion to be met, the statistic of the interaction between listen and turn number had to be positive (slope in Listen 1 > slope in Listen 2) and significant at $q_{FDR} < 0.05$. This was our most important criteria– see *Combining these criteria.*

We additionally expected the slope of Listen 2 to be positive. We tested this by fitting a LMEM for each region, predicting the turn-template pattern correlation (Fisher z-transformed) in Listen 1 from the speech turn number (with higher numbers corresponding to later speech turns) and treating participant as a random effect. For this criterion to be met, the estimate had to be positive.

*Supporting analysis: pre- versus post-twist updating rates.*

To aid comparison with the narrative- and episode-level analyses and to further clarify the temporal dynamics underlying character stabilization, we conducted an additional supporting analysis examining whether the rate of character updating differed between the pre- and post-twist segments of the narrative. For each participant and region, we estimated the linear relationship between event order and Fisher z–transformed event–template correlation separately for pre- and post-twist speech turns, treating character as a random effect. We then compared the resulting slope estimates using paired t-tests across participants. This analysis probes whether character representations update more rapidly during the pre-twist portion of the narrative, when character identity is still uncertain, than during the post-twist portion, when representations are closer to asymptote and thus more stable. As expected, slopes were steeper during the pre-twist segment in Listen 1.

To confirm that this pre–post asymmetry was specific to initial exposure to the narrative rather than reflecting a generic early–late effect, we applied the same analysis in Listen 2 and compared the pre–post slope differences across listens. Consistent with the primary stabilization analysis, the pre–post slope difference was reduced in Listen 2, reflecting diminished online character updating once the twist was known. In both cases (pre > post in L1 and $L1_{pre>post} > L2_{pre>post}$), the t-value had to be positive and significant at $q_{FDR} < 0.05$.

*Criterion 4—Control: Representations of the two characters are distinct from one another in Listen 1 and Listen 2.*

As a control, we compared representations of the other character to the final template of the character being tested (e.g., Steve turns relative to Lucy's template, and vice versa). Specifically, we computed turn-template correlations using the other character's speech turns from Listen 1 and Listen 2. We then fit two independent LMEMs for each listen, with random effects for participant and for template character (Lucy vs. Steve), predicting this Fisher z–transformed correlation from the turn number. Given that we expected representations of the other character to be distinct, we did not expect a positive correlation with the template. Therefore, for this criterion to be met, the estimates for each of the models needed to be negative or at least not significantly positive. This would indicate that the representations of the characters are becoming increasingly distinct.

A summary of the criteria and their corresponding representation in the schematic in **Fig. 4B** is below. As noted, all correlations were Fisher z-transformed.

- **Criterion 1** (solid red line in **Fig. 4B** schematic): the slope of same-character turn-template correlations over time had to be positive.
- **Criterion 2** (arrow pointing to 'turn 1'): the correlation between the first character speech turn and the template had to be positive in Listen 2 and greater than the same coefficient in Listen 1.
- **Criterion 3** (dashed red-line relative to solid red line): the slope of Listen 1 had to be significantly greater than the slope during Listen 2, but the slope during Listen 2 still had to be positive.
- **Criterion 4** (negative gray lines): correlations of speech turns with the other character's template had to be negative or not significantly positive.

*Combining criteria.*

We had two steps to combining these criteria. First, to be conservative, we show only regions that fit all of the expected criteria (no effect in Criterion 4 and positive in Criteria 1-3) in our character representation maps (**Fig. 4C**). Second, we considered Criterion 3 – *Representations of the character stabilizes in Listen 2* – as the most critical, given its focus on within-subject, across-listen updating and reloading of character representations. Consequently, we used estimates from this analysis for plotting and for subsequent analyses (see *Clustering results across analyses*). We also used the corresponding *p*-values to correct for multiple comparisons using FDR with an alpha of 0.05 across the 97 regions for this criterion.

*Testing template reliability.*
We additionally tested the consistency and reliability of our character templates. First, we examined template stability across three different template sizes. **Supplementary Fig. 4B** shows similar outcomes for both characters when using longer templates (i.e., created by taking the median across the last three or four speech turns, versus the last two speech turns used in the main analysis).

Second, we tested template consistency using split-half reliability analysis. We hypothesized that given the updated narrative model within the post-twist segment, the template would remain relatively stable regardless of which specific turns were used, as long as they came from this segment. For each brain region and character, we calculated split-half reliability by randomly dividing trials into two halves, computing median spatial patterns for each half, and correlating these patterns across 100 iterations per subject. Correlations were Fisher z-transformed before analysis. Individual-level significance was assessed via permutation testing (100 permutations) against spatially shuffled null data. Group-level significance was tested using one-sample t-tests against zero.

For Lucy, across the 45 regions showing effects in our main analysis (**Fig. 4C**), an average of 35.9 ± 0.4 participants (~100 %) per region demonstrated individually significant split-half reliability ($p < 0.05$). At the group level, 75.6% (34/45) of regions showed significant reliability. Most participants showed positive split-half correlations as expected (mean proportion = 0.655 ± 0.080, range = 0.500 to 0.806), where 1.0 indicates all participants in a region show positive reliability and 0.5 indicates an even split between positive and negative correlations. For Steve, across the 45 regions from the main analysis, an average of 35.6 ± 0.6 participants (~99%) per region showed individually significant reliability. Group-level significance was observed in 15.6% (7/45) of regions, with slightly fewer participants showing positive split-half correlations (mean proportion = 0.521 ± 0.093, range = 0.333 to 0.694).

The somewhat lower consistency for Steve likely reflects reduced statistical power from fewer post-twist speech turns (9 vs 16 turns). Nevertheless, these results demonstrate that most character-responsive regions show reliable and consistent activation patterns in the majority of subjects, providing strong validation for the stability of our template approach.

**Clustering results across analyses.**
In our final analysis, we aimed to assess the extent to which representations of the three narrative elements—narrative models (**Fig. 2D**), episodes (**Fig. 3C**), and characters (**Fig. 4C**)—rely on overlapping versus distinct brain regions. To compare effect estimates across the three elements, we first set estimates for any region that did not fit the expected direction for a given element to 0. (For the character analysis, regions had to fit the expected direction in all four of our criteria.) This resulted in 34 regions being set to 0 in characters, 22 in episodes, and 2 in the narrative model. We then normalized the estimates across regions within each element analysis using min-max scaling; the scaled values allowed for consistent comparison across analyses.

We took two approaches to comparing region-wise involvement across the three narrative elements. First, we correlated the scaled estimates across regions between all possible pairs of the three analyses. Second, we used KMeans clustering [19] to perform pattern vector-based clustering, grouping regions based on the similarity of their pattern of estimates derived from each analysis.

This approach captured the underlying structure of the relationships across regions and assigned a single cluster label to each region. To determine the optimal number of clusters ($k$), we first calculated the silhouette score for values of $k$ ranging from 2 to 10 (see **Supplementary Fig. 5A**).

We selected $k = 4$ because it yielded both a high silhouette score ($s = 0.43$) and some amount of dissociation between elements; a $k = 2$ solution had a higher score ($s$=0.46), but simply grouped all regions responsive to *any* of the three elements together in one cluster with the second cluster consisting of the regions largely uninvolved in any level of representation (see **Supplementary Fig. 5B).**

**Visualization.**
Motivated by a recent recommendation [20], we present largely unthresholded, whole-brain maps for all of our main figures and add contours to indicate regions meeting criteria for statistical significance (described in detail in the caption of each figure). Although our discussion mostly focuses on only those regions meeting statistical significance, we display results across the brain to provide insight into the directionality of effects and facilitate comparisons with past and future work.

To do so, we create a translucent map weighting the data by an opacity (alpha) mask using a threshold of 20% of the maximum range of the data. For each region, we calculate an alpha value (ranging from 0 to 1) to determine its transparency level. If a value exceeds a threshold of 20%, the corresponding region is assigned an alpha value of 1 (fully opaque) and is normally plotted. For values between 0 and our threshold, the alpha value is scaled proportionally from 0 to 1, with increasing transparency for weaker effects. In the narrative model and episode analyses, regions meeting a corrected threshold of $q_{FDR} < 0.05$ are outlined in black. In the character analysis, all plotted regions meet the criteria at their respective thresholds, with contours indicating whether a region satisfies the criteria in both characters (in dark blue). The SurfPlot package [21,22] was used for visualization.

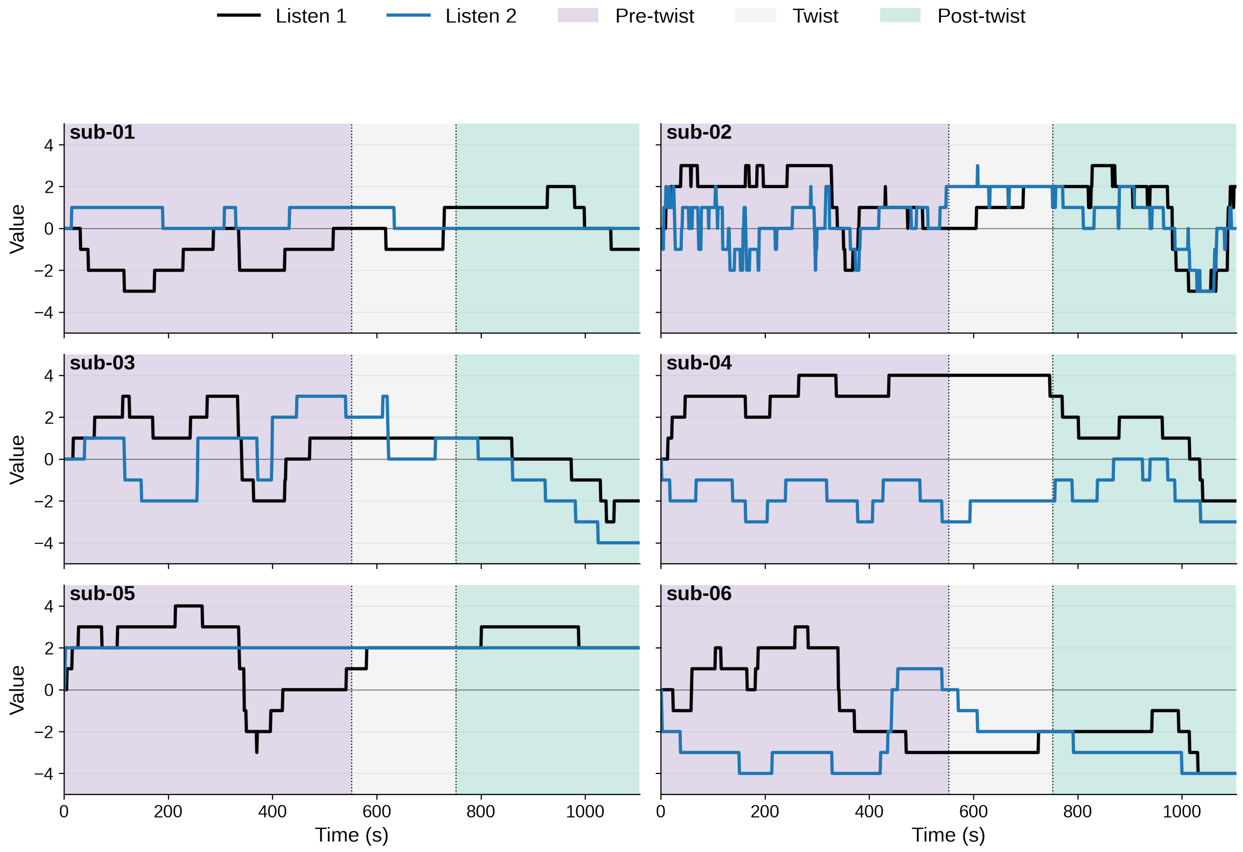


***Supplementary Fig.1.*** Sample behavioral timeseries. Sample behavioral timeseries from the first 6 subjects are shown to highlight the degree of variability in slider usage.

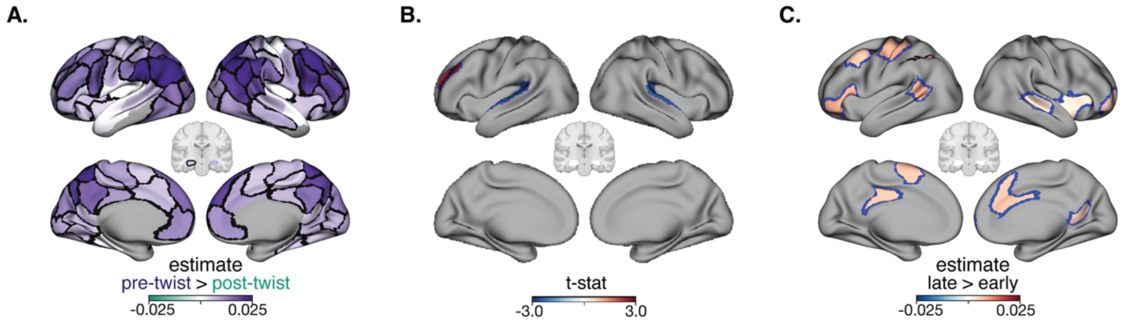


***Supplementary Fig.2. Narrative model control analyses.*** **A. Pre- versus post-twist differences do not change with continuous rating button presses regressed out.** Per individual participant, we regressed the timeseries of their button presses in the continuous rating task from their BOLD signal timeseries and reperformed our analyses on the residuals of this regression. Results are largely unchanged (compare to **Fig. 2D**). Regions contoured in black show an FDR-corrected significant effect at $q_{FDR} < 0.05$ for all matched-length comparisons between neural shifts pre- and post-twist. **B. Pattern intra-SC does not generally increase over time in an independent stimulus.** Using repeated listens of an auditory story from LeBel *et al.* (2023), which did not have a twist or induce any substantial interpretation change, we modeled pattern intra-SC as a function of time. The only regions that show a significant effect of time ($p < 0.05$, uncorrected) on within-subject pattern correlations are bilateral auditory cortex (negative effect such that participants become less similar to themselves over time; blue) and the left dorsolateral PFC (positive effect such that participants become more similar to themselves over time; red). **C. Pattern intra-SC is slightly higher within the later half of a narrative segment, but within-segment effects are weaker than across-segment effects.** We divided both the pre-twist and post-twist segments into "early" and "late" halves and compared the median intra-SC values, per individual, across these timepoints, analogous to our main analysis comparing across segments (pre- versus post-twist). Using a linear mixed effects model with random effects of subject and fixed effects of segment and half (early versus late), we tested the hypothesis that the later half of a segment would show greater median pattern intra-SC than the earlier half. Estimates plotted show the strength of the difference between segment halves. Regions contoured in black show an effect at $q < 0.05$ (corrected for multiple comparisons using the false-discovery rate). Regions contoured in blue show an effect at $p < 0.05$ (uncorrected). The only region to show effects at $q < 0.05$ was the left intraparietal sulcus (IPS). Although few of the same regions (n=4: left posterior cingulate, left frontal eye fields, right vmPFC, and the left IPS), emerged as in our main analysis, as hypothesized, late versus early effects were weaker within segments than across segments (compare to **Fig. 2D**).

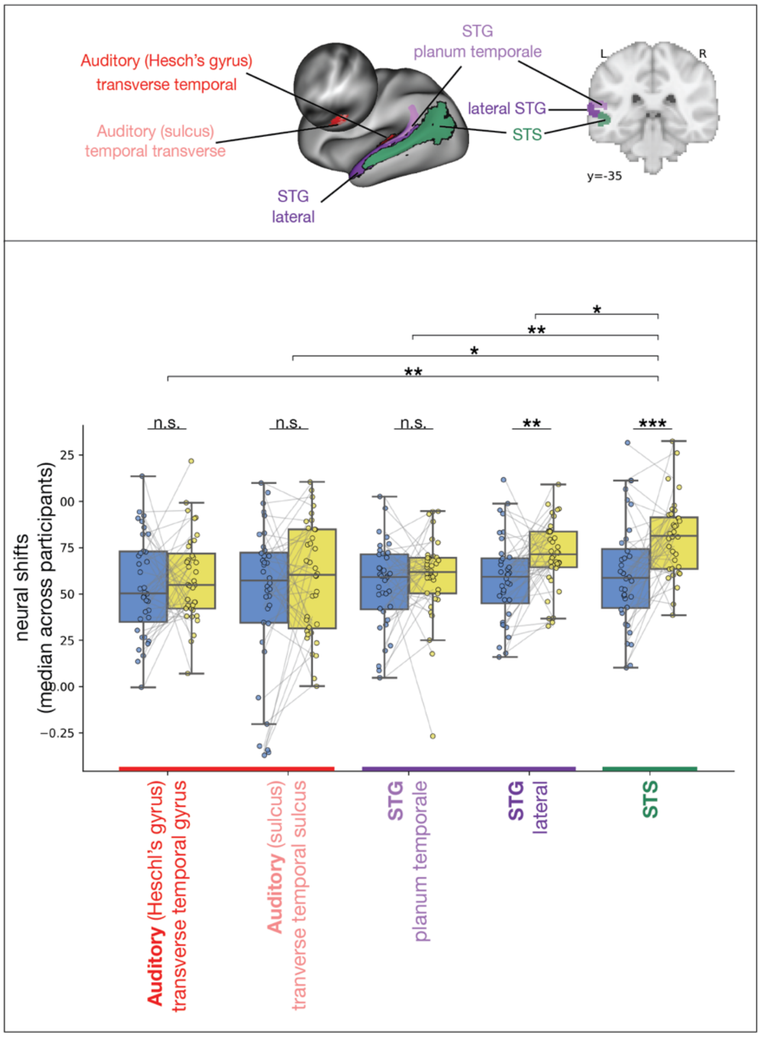


***Supplementary Fig.3. Left superior temporal sulcus shows stronger episode effects than neighboring temporal regions.*** Neural shifts for reevaluated versus control episodes were compared across left temporal lobe regions, including auditory cortex (red-tinted regions), subdivided into the transverse temporal gyrus (Heschl's gyrus) and the transverse temporal sulcus; the superior temporal gyrus, STG; purple), subdivided into the planum temporale and lateral STG; and the superior temporal sulcus (STG; green).

Blue boxplots represent control episodes, and yellow boxplots represent reevaluated episodes. Estimates reflect the strength of the difference between reevaluated and control episodes within participants. These estimates were derived from a linear mixed-effects model in which within-subject neural shifts were predicted by episode type (coded as reevaluated > control), with participant and episode pair included as random effects. Dots indicate each participant's median neural shift across episodes of each type. Statistical significance markers reflect FDR-corrected p-values (* $q < 0.05$, **$q < 0.01$, ***$q < 0.001$). Although significant effects were also observed in lateral STG (dark purple; $\beta = 0.072$, $q = 0.002$), the STS exhibited by far the largest effect (green; $\beta = 0.10$, $q < 0.0001$). Further, differences between reevaluated and control episodes were compared for each region statistically, highlighting again that the strongest effects are in the STS. The STS showed significantly stronger effects than every other temporal region examined, including Heschl's gyrus ($t = 3.57$, $q < 0.01$), the transverse temporal sulcus ($t = 2.57$, $q < 0.05$), the planum temporale ($t$ = 3.98, $q$ < 0.01) and lateral STG ($t$ = 2.40, $q$ < 0.05). Not visualized: while the STG as a whole showed weaker effects than the STS, the lateral portion of the STG nonetheless exhibited significantly stronger effects than both the planum temporale (light purple; $t$ = 2.49, $q$ < 0.05) and Heschl's gyrus ($t$ = 2.71, $q$ < 0.05).

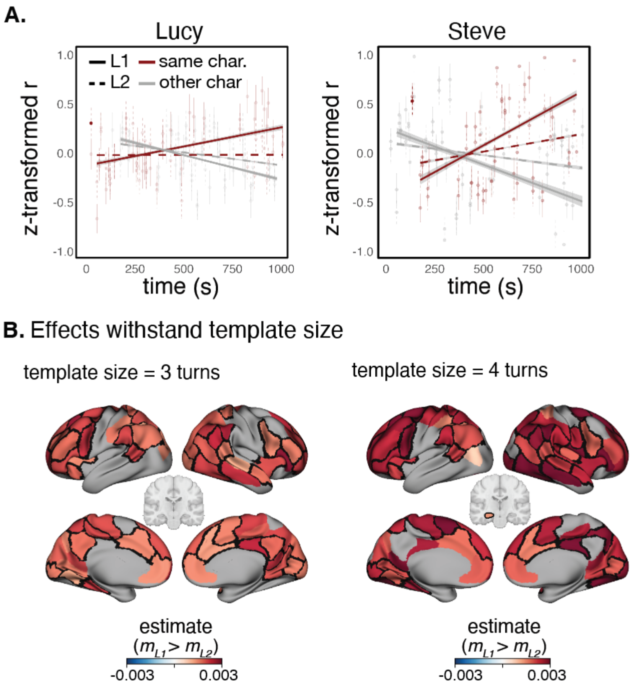


***Supplementary Fig.4. Character templates are robust*.** **A. Template correlations in a sample region.** Data for the left TPJ for the Lucy and Steve template analyses (**Fig. 4C**). Fisher Z-transformed r-values between each event and the template are plotted on the y-axis. For visualization, linear trend lines (solid/dashed lines) are fitted to predictions from a mixed-effects model (r ~ event × listen × character + (1|participant)) run per parcel. Lines show trends by listening condition (L1 = solid, L2 = dashed) and character (red = same-character [e.g., Lucy-Lucy], gray = other-character [e.g., Lucy-Steve]). Events were matched to their onset times in the story, accounting for variability in line start times. Error bars represent standard error of the mean from raw (z-scored) subject data. The first L2 event is highlighted with increased opacity to mark condition onset. Despite variability in strength, the data fit the hypothesized relationships shown in the schematic in **Fig. 4B**. **B. Effects withstand template size.** Templates of size 3 and 4 speech turns were tested. Compare panels to **Fig. 4C**. Generally, the same regions show effects regardless of template size. Estimates are shown for regions that show an effect in the expected direction for all criteria. Contours reflect significance in the main criterion at $q_{FDR}$ < 0.05.

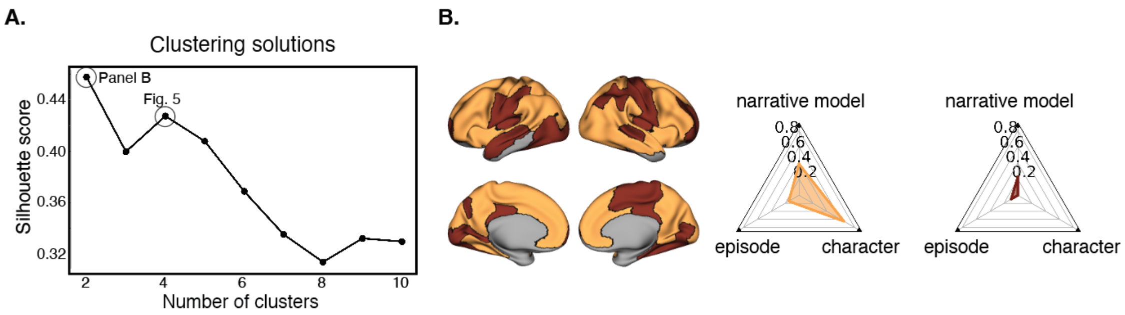


***Supplementary Fig.5.*** **Clustering solution. A.** The silhouette score across different numbers of clusters is shown. **B.** A solution of $k$ = 2 is shown.

***Supplementary Table 1: Sample recall transcripts.*** Transcripts were minimally edited to remove disfluencies (e.g., 'like', 'kind of') and for readability, while preserving original meaning. The data from the first three participants is shown. These were not chosen for specific content, but rather to be representative of participants' speech looked like generally. Instructions can be found in the main text and are copied here: After Listen 1, they were instructed "*Using the microphone, please describe the moments at the beginning of the story that you reconsidered after hearing the end.*" After Listen 2, they were instructed, "*Using the microphone, please describe the moments of the story that changed for you after hearing the story once before.*"

| Subject Number | Listen | Transcript |
|---|---|---|
| 1 | L1 | I remember it feeling like it was in the 1950s and it did not seem like at the end that we were in the 1950s. It was kind of in a post-apocalyptic world in the 2050s. And I assumed Lucy was kind of popular... She seemed like a little ditzy woman who was concerned about following all of the rules of the store. But later, it was revealed that she was a robot who was designed to maintain the status quo in the face of an apocalypse. So, I reframed how I saw her based on the information that she was programmed rather than had free will. |
| 1 | L2 | I think I missed more or less what happened at the end of the world the first time. So, this time I noticed that it was nanotechnology and that there were kind of these, like, mutant human things that were roaming around. I also noticed smaller details, especially in the beginning of the story, when Lucy was describing the dog roaming around and aspects of the mall setting. I noticed that more starkly the second time listening [to the story]. And yeah, I just kind of filled in a little bit more of the more granular details this time around. |
| 2 | L1 | So, at the beginning of the story, we hear Lucy talking to her boyfriend on the phone and you later realize that this boyfriend is just a subroutine that she is playing in her mind, that she's programmed to rehearse because she's a robot. I also reconsidered the dog that she hears running around that she keeps calling security for. The dog—actually what she thinks is a dog—turns out to be a mutated human that's hungry and looking for something to eat. And, at first, I thought Steve was really going to be a bit of a jerk or a homeless person. But it actually turns out that he's someone who's trying to escape from these mutated demons and persecution. |
| 2 | L2 | So, something that changed [from] when I heard the story before…one thing that I just picked up at the very end was her hearing that her battery was at 7%, which kind of confirmed that she was a robot. The vague conversation with her boyfriend, I definitely found more humorous, because you get this idea that she's really just running through the subroutine. In general, I found the things that were clearly subroutines or robot-isms to be more humorous the second time, even though it was a little bit more frustrating to hear her not help Steve because there was no ambiguity about her being a robot. |
| 3 | L1 | At the beginning of the story, she was talking to her boyfriend about watching the game and making out with him. But after hearing what Steve said about how the robots have fake families and lives at home and she called him, it made me reconsider who she was talking to and whether or not it was due to her imagination or if she actually had a boyfriend or if it was really fake. And that at the beginning she hears the dog barking and describes that to her boyfriend, and then when the dog stops barking, [I] assumed that the dog had eaten Steve and that was what had fed him in her mind. Then she describes in her fake-robot-mind boyfriend that the dog stopped barking, and she couldn't hear it anymore so it must have left the mall. |
| 3 | L2 | It was interesting to hear how Steve's part of the story changed so much because a lot of the things he said when he called her a robot in the beginning, it just sounded like he was mad at her for not helping him. But in |

| | | |
|---|---|---|
| | | reality, he was actually calling her a robot. And then she's talking about the music and how that's her favorite song on the radio. And when he's talking about the food court and the rubble [and] how you can't get water from there, even though she's asking him to. And that the boyfriend at the beginning [when they are] talking about the game and then when [she] brings up the game later [saying] how a robot will be programmed to talk about the big game and have a boyfriend... even though he doesn't actually exist and is just a part of her robot brain. |
